# Machine perfusion and single-cell spatial transcriptome mapping identifies novel immune escape mechanisms in colorectal cancer liver metastasis

**DOI:** 10.64898/2025.12.09.693114

**Authors:** Peter Kok-Ting Wan, Ranchu Cheng, David Johnson, Carl Lee, Shihong Wu, Areeb Mian, Ahmet Hazini, Sorayya Moradi, Kate Friesen, Flurin Caviezel, Syed Hussain Abbas, Hatem Sadik, Adil Lakha, Keaton Jones, Girishkumar Kumaran, Matthew Bottomley, Rob Jones, Mark Coles, Rachael Bashford-Rogers, Constantin Coussios, Len Seymour, Robert Carlisle, Kerry Fisher, Alex Gordon-Weeks

## Abstract

Liver metastasis is the terminal stage of colorectal cancer. Immune checkpoint blockade (ICB) has heralded remarkable clinical success across a range of cancer types, but with limited efficacy in replacement-type colorectal liver metastasis (CRLM), the most common and lethal histological subtype. Using a novel human normothermic perfusion model, we demonstrate for the first time, that T-cells preferentially extravasate within the peri-tumoural liver rather than the CRLM. We use single-cell spatial transcriptomics and multiplexed immunofluorescence to validate CRLM T-cell exclusion and show that CRLM endothelia are anergic, lacking key receptors required for T-cell extravasation. CD4 T-cells that extravasate within the peri-tumoural liver are TCR-reactive, yet exhausted, whilst CD4 T-cells within the CRLM demonstrate a stress response, impaired cytokine expression and lack of TCR reactivity. We identify the spatial cellular and molecular interactions underlying these observations, providing novel targets for future attempts to sensitise to immunotherapeutics and a justification for failed ICB efficacy in CRLM.

## Introduction

Liver metastasis is the most common cause of colorectal cancer death and is diagnosed with increasing frequency, presenting a pressing clinical need^1^. The liver is a fertile metastatic soil due to a massively parallelled sinusoidal structure, high blood flow, and tolerogenic microenvironment^2^. By the time they are clinically apparent, CRLM achieve an advanced evolutionary state, through interaction between mutational and epigenetic events and selective pressures afforded by the various tissue environments of the colon, blood, lymphatics and liver^3,4,^. A major component of metastatic evolution is immune evasion^5,6^, which serves as a barrier to successful CRLM treatment.

Several studies have investigated CRLM immune evasion using sc-RNAseq^7,8,9,10,11,12^, yet sample disaggregation generates small cell numbers, loss of rare, fragile populations and inability to understand spatial context and structure; features that have provided biological insights in CRC^13,14^. This has hampered understanding of how the CRLM co-opts the liver to gain and maintain a survival advantage. Spatial resolution is particularly important for investigating the biologically important region between the CRLM and the liver, which cannot be adequately profiled using disaggregated tissue. Although partially addressed in spatial transcriptomic studies^12,15^, until recently^16^, these were not at single cell level, requiring deconvolution for cell composition inference.

We address these shortcomings by combining a human whole tumour CRLM perfusion platform with single-cell protein-based and transcriptomic analysis of large areas of CRLM and background liver, using multiscale spatial techniques (single cell, neighbourhood, topological). We demonstrate that CRLM utilise endothelial anergy and CD4 T-cell dysfunction as primary mechanisms of immune escape.

## Methods

### Tissues and ethics

All tissues were retrieved from patients undergoing surgical resection of liver metastatic colorectal cancer within the Department of Hepatobiliary Surgery, Oxford, UK and Department of Hepatobiliary Surgery, Liverpool, UK. Tissues were collected under ethics numbers 22/SC/0429 and 21/YH/0206. The clinical details of patients in the CosMx and Cell DIVE datasets and those in hemi-liver perfusion experiments are shown in supplementary tables 1-3 respectively.

### Testing T-cell extravasation in-vivo

#### Normothermic liver perfusion

Hemi-livers containing CRLM were selected based on favourable hepatic inflow and outflow vascular anatomy on review of triple phase CT and MRI by the operating surgeon (Gordon-Weeks). Surgical resection proceeded in a way as to limit warm ischaemia to <10 minutes, through hepatic parenchymal transection first, with transection of inflow and outflow vasculature immediately before specimen explant. Extra-hepatic vascular length was maintained to aid cannulation. Inflow clamping (Pringle manoeuvre) was applied for 10-minute intervals intra-operatively with 5-minute breaks to limit reperfusion injury. Hemi-livers were flushed immediately on explant with 1L ice-cold Custodiol® perfusion fluid containing 10,000 IU of unfractionated heparin (UFH) (5 IU/mL, Panpharma, Luitré, France) on crushed ice, followed by further Custodial® until the liver blanched completely. OrganOx *metra*® inflow/outflow cannula were secured. Meanwhile, the *metra*® was primed with 3 units of O Rh negative packed red blood cells (Cambridge Biosciences, Cambridge, UK), with 500-1000 mL Gelofusine, cycled through the device for pO2 and pCO2 conditioning until optimal temperature (37 °C) and perfusate pH (7.3-7.4) were reached through sodium bicarbonate administration (B. Braun Medical ltd., Hessen, Germany). Meropenem (500 mg, Consilient Health, Dublin, Ireland) and 10,000 IU heparin were added to the perfusate.

The hemi-liver was onboarded and perfusion started. Throughout perfusion, the following were infused; 0.5 g epoprostanol sodium (Flolan, MercuryPharma, London, UK), 8 μg/hr, 25,000 IU heparin, 833 IU/hr, 5.6 g sodium taurocholate (OrganOx, Oxford, UK), 187 mg/hr, and 200 IU fast-acting insulin (Actrapid, Novo Nordisk, Bagsvaerd, Denmark) titrated to perfusate glucose concentration. Nutriflex parenteral nutrition solution (B. Braun Medical ltd., Kronberg, Germany) was introduced, dependent on glucose concentrations. Arterial and venous flows were regulated autonomously by the *metra*® platform and integrated in-line arterial blood monitoring (Terumo CDI 550; Terumo, 58 Tokyo, Japan) provided O_2_ and CO_2_ partial pressure and pH readings per second. Arterial blood gas measurements, haematocrit and haemoglobin levels were recorded 4-hourly (i-STAT 1, Abbott, Illinois, USA). Liver biochemistry was measured 8-hourly (Piccolo Xpress, Abbott, Illinois, USA).

At experimental termination, livers were perfused with a further 3L Custodiol® to remove circulating lymphocytes, before perfusion with 500mls Lycopersicon Esculentum (Tomato Lectin, Sigma Aldrich) diluted in saline to a final concentration of 2μg/ml.

### PBMC isolation

PBMCs were obtained from ABO-compatible whole-blood leukocyte cones (NHS Blood and Transfusion Service) using SepMate PBMC isolation tubes (STEMCELL Technologies, UK). PBMC’s were subjected to red blood cell lysis buffer (Qiagen, Germany) and washed with DPBS. The PBMC pellet was resuspended in 2% RPMI supplemented with 2% FBS and stained with CellTrace™ CFSE Cell Proliferation Kit (Invitrogen, UK). PBMCs were washed with DPBS and resuspended in DPBS on ice for injection.

### FACS analysis

Liver and tumour excisional biopsies were dissociated (Tumour Dissociation Kit, Miltenyi Biotech, UK). The resulting cell suspension was strained (70µM) and washed with MACS buffer. Whole blood samples were centrifuged and treated with RBC lysis buffer followed by washing with MACS buffer. Cells were stained with Live/Dead Fixable Near-IR dead cell stain (Thermofisher Scientific, UK) and fixed with 10% neutral buffered formalin (Merck). Antibodies used for cell-type detection are shown in table 2. Flow cytometric data acquisition was performed on Cytoflex Flow Cytometer (Beckman Coulter, UK) and analysed using FlowJo v10.10.0 software (BD Biosciences).

**Table 1.**
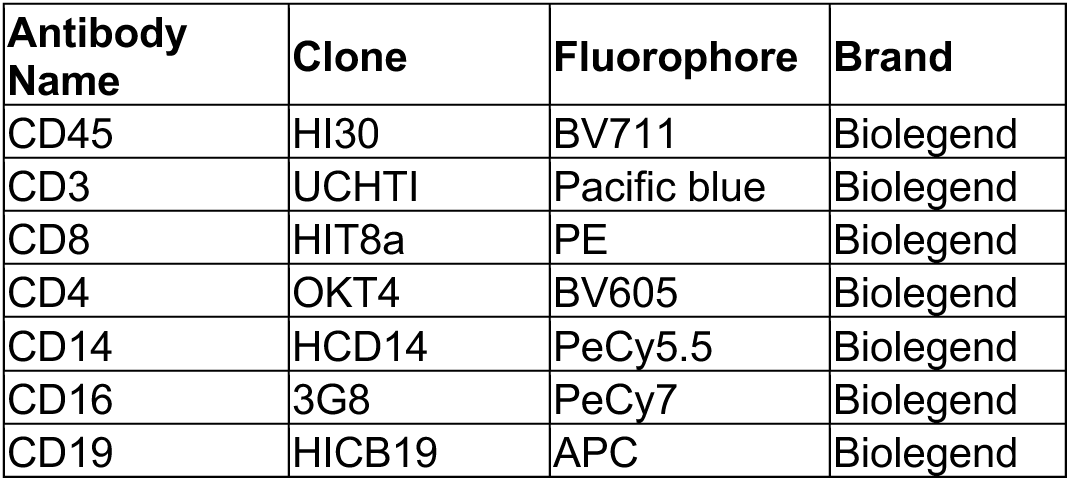
Antibodies used in FACS experiments.

| **Antibody Name** | **Clone** | **Fluorophore** | **Brand** |
| --- | --- | --- | --- |
| CD45 | HI30 | BV711 | Biolegend |
| CD3 | UCHT1 | Pacific blue | Biolegend |
| CD8 | HIT8a | PE | Biolegend |
| CD4 | OKT4 | BV605 | Biolegend |
| CD14 | HCD14 | PeCy5.5 | Biolegend |
| CD16 | 3G8 | PeCy7 | Biolegend |
| CD19 | HICB19 | APC | Biolegend |

**Table 2.**
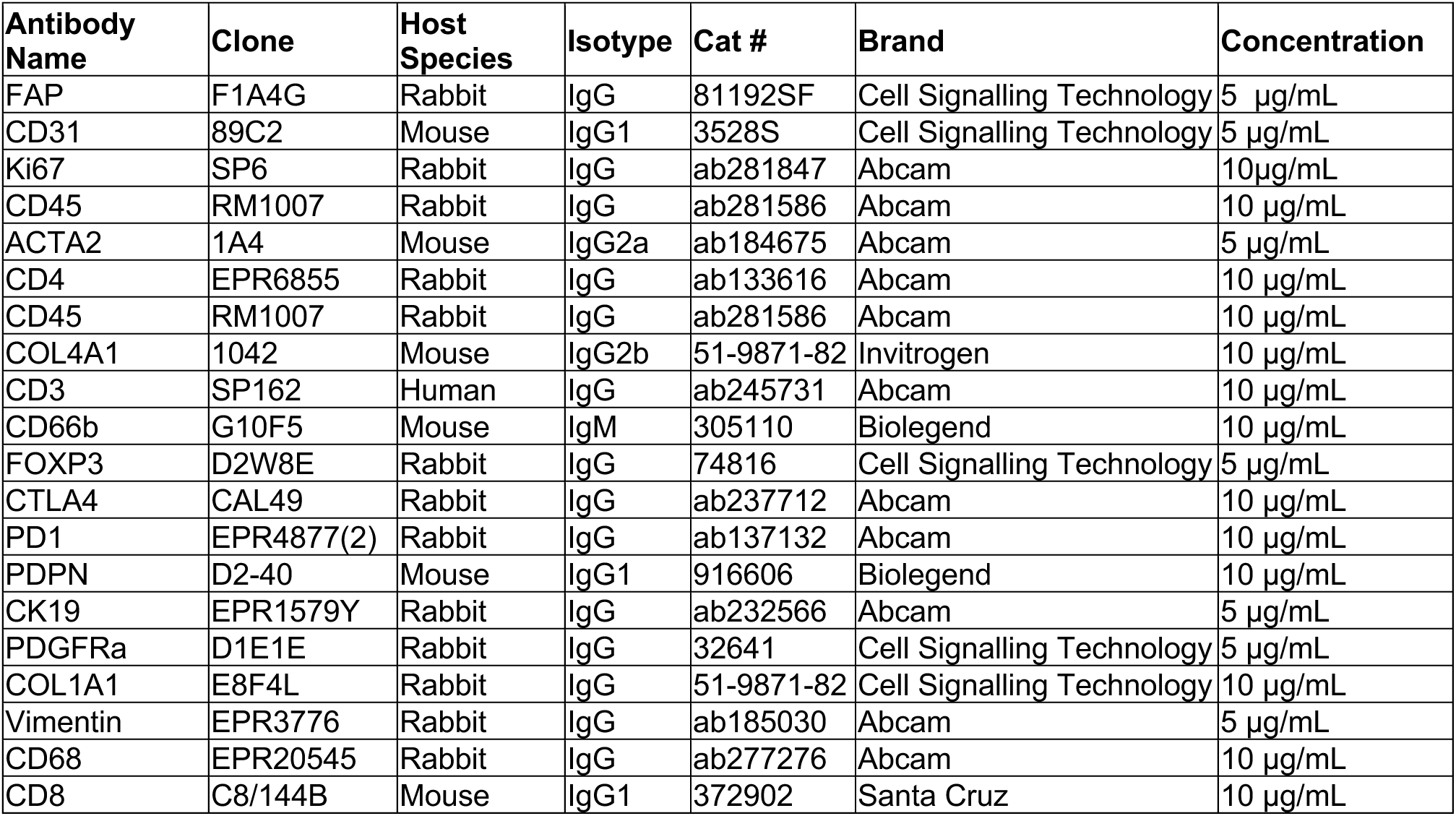
Antibodies used in Cell-DIVE mIF.

| **Antibody Name** | **Clone** | **Host Species** | **Isotype** | **Cat #** | **Brand** | **Concentration** |
| --- | --- | --- | --- | --- | --- | --- |
| FAP | F1A4G | Rabbit | IgG | 81192SF | Cell Signalling Technology | 5 µg/mL |
| CD31 | 89C2 | Mouse | IgG1 | 3528S | Cell Signalling Technology | 5 µg/mL |
| Ki67 | SP6 | Rabbit | IgG | ab281847 | Abcam | 10 µg/mL |
| CD45 | RM1007 | Rabbit | IgG | ab281586 | Abcam | 10 µg/mL |
| ACTA2 | 1A4 | Mouse | IgG2a | ab184675 | Abcam | 5 µg/mL |
| CD4 | EPR6855 | Rabbit | IgG | ab133616 | Abcam | 10 µg/mL |
| CD45 | RM1007 | Rabbit | IgG | ab281586 | Abcam | 10 µg/mL |
| COL4A1 | 1042 | Mouse | IgG2b | 51-9871-82 | Invitrogen | 10 µg/mL |
| CD3 | SP162 | Human | IgG | ab245731 | Abcam | 10 µg/mL |
| CD66b | G10F5 | Mouse | IgM | 305110 | Biolegend | 10 µg/mL |
| FOXP3 | D2W8E | Rabbit | IgG | 74816 | Cell Signalling Technology | 5 µg/mL |
| CTLA4 | CAL49 | Rabbit | IgG | ab237712 | Abcam | 10 µg/mL |
| PD1 | EPR4877(2) | Rabbit | IgG | ab137132 | Abcam | 10 µg/mL |
| PDPN | D2-40 | Mouse | IgG1 | 916606 | Biolegend | 10 µg/mL |
| CK19 | EPR1579Y | Rabbit | IgG | ab232566 | Abcam | 5 µg/mL |
| PDGFRa | D1E1E | Rabbit | IgG | 32641 | Cell Signalling Technology | 5 µg/mL |
| COL1A1 | E8F4L | Rabbit | IgG | 51-9871-82 | Cell Signalling Technology | 10 µg/mL |
| Vimentin | EPR3776 | Rabbit | IgG | ab185030 | Abcam | 5 µg/mL |
| CD68 | EPR20545 | Rabbit | IgG | ab277276 | Abcam | 10 µg/mL |
| CD8 | C8/144B | Mouse | IgG1 | 372902 | Santa Cruz | 10 µg/mL |

### Protein Imaging, QC and segmentation

Multiplexed immunofluorescence (mIF) was performed on formalin-fixed, paraffin-embedded (FFPE) tissue sections using the Cell DIVE™ platform (2500HS, Leica Microsystems, Germany). Following deparaffinization and a two-step antigen retrieval (citrate buffer pH 6.0 then Tris-EDTA buffer pH 9.0), slides were permeabilized with 0.05% Triton X-100, then blocked with 1% BSA, 1% normal goat serum and 1% normal donkey serum in PBS containing 0.05% Tween-20. Slides were initially stained with DAPI and imaged across all fluorescence channels to capture tissue-specific autofluorescence. In the first staining round, slides were incubated with three antibodies (table 2) and imaged. All rounds employed directly conjugated primary antibodies targeting up to three markers per cycle, with DAPI included in each round. Whole-slide images were acquired using a motorized widefield fluorescence microscope equipped with a 20×/0.75 NA objective. Fluorescence was collected sequentially in DAPI (∼460 nm), FITC (∼520 nm), Cy3 (∼570 nm) and Cy5 (∼670 nm) channels. Exposure times were optimized using single-stained controls and held constant across all samples. Between cycles, fluorophores were inactivated using three consecutive 15-minute incubations in fresh alkaline hydrogen peroxide buffer (3% H₂O₂ in 20 mM NaHCO₃, pH 11.2), protected from light. Slides were re-imaged to confirm complete signal removal and capture new autofluorescence profiles, then de-coverslipped for the next staining cycle. This stain–image–bleach sequence was repeated until the full panel was completed. Image stacks from all cycles were computationally registered using DAPI nuclear features, and downstream analysis was performed using the Cell DIVE image processing pipeline.

### Cell-DIVE image QC, segmentation and cell phenotyping

Raw imaging tiles were stitched and merged using ASHLAR^60^ to yield composite images. Autofluorescence was performed using the ‘background_subtraction’ module (MCMICRO)^61^. Single-cell segmentation was conducted using Mesmer^62^ using nuclear staining and a composite membrane/cytoplasmic signal generated by concatenation of multiple stains (CK19, αSMA, CD31, CD3, CD4, CD8, CD68). Quantification of per-cell marker expression was performed using the marker_quantification module in the ark-analysis framework^63^.

Phenotyping was performed manually. Arcsinh-transformed marker intensity values derived from the whole-cell segmentation were used as input for gating. All markers were visually inspected, and gating thresholds independently determined per marker and for each image using the SCIMAP gate_finder tool^64^. Following threshold determination, expression values were rescaled, and markers with normalized intensities > 0.5 were considered positive. Cell type annotation was performed using a hierarchical phenotyping workflow implemented via the SCIMAP phenotype_cells function^64^. Accuracy was assessed through iterative visual overlays of phenotypes onto the original multi-channel images, with additional refinement performed when discrepancies between marker expression and spatial localization were observed.

Segmentation errors and imaging artifacts were systematically identified and excluded. Cells with a nuclear-to-cell ratio greater than 1 were flagged as abnormal and removed. For every imaging cycle, marker co-expression patterns were assessed to detect signal anomalies. In cases where mutually exclusive markers showed abnormal strong co-localized signals within the same region, intensity thresholds were determined, and affected cell were labelled as double-positive artifacts and subsequently excluded. For non-exclusive markers presenting unusually bright signals or patterns suggestive of imaging artifacts, thresholding was performed manually with expert input and visual check to ensure accuracy. The scripts needed for the preprocessing above are openly available at: https://github.com/ShihongWu/multiplexed-image-preprocessing.git and described in detail in SpatioEv (doi: https://doi.org/10.1101/2025.06.30.662328).

### Quantification of CD3 counts in portal triad regions

Image segmentation and analysis were performed in QuPath. Cell-DIVE images were limited to markers of the portal triad including Col4a1, CD31 and CK19 enabling positive identification of portal triads in the background and peri-tumoural liver. To define the portal triad as a separate ROI, we triangulated the most distant visible triad structures (bile duct, hepatic artery or portal vein). Example of resultant ROIs are demonstrated in Supplementary Figure 6B. CD3-positive cell count was normalised against total cell count within each corresponding ROI and presented as a percentage.

### CosMx imaging, QC, cell segmentation and phenotyping

Tissues were prepared and processed on the CosMx Spatial Molecular Imager (Bruker Spatial Biology, USA) as per the CosMx SMI Manual Slide Preparation for RNA Assays and CosMx SMI Instrument User Manual. 5 μm thin FFPE sections were mounted to Leica Bond Plus slides (Leica Biosystems, Richmond, USA) within 2 weeks of imaging and baked overnight at 60°C to increase tissue adherence. After baking, the slides were dewaxed and antigen retrieval carried out using Target Retrieval Solution (Bruker Spatial Biology, USA). Tissues were permeabilised using 3 μg/mL Proteinase K and fiducials applied, followed by post-fixation in 10% neutral buffered formalin (NBF). Tissues were washed with NBF stop buffer and then treated with fresh acetate solution and the slides hybridised with CosMx Human 6K probe mix and incubated overnight^65^.

The slides were then washed with stringent buffer in a water bath twice before incubation with in blocking buffer. This is followed by morphology and segmentation marker staining using CD298/B2M mix (Bruker Spatial Biology, USA) and PanCK/CD45 mix (Bruker Spatial Biology, USA) as recommended in the standard protocol for non-neuronal tissues. Following this, flow cells were prepared using CosMx Flow Cell Assembly Kit and loaded on to the CosMx Spatial Molecular Imager. Configuration C was used for Pre-Bleaching profile, and Configuration E was used for Cell Segmentation profile. Following image acquisition, FOVs were selected and cycling and data collection were initiated. FOVs were specifically selected to cover regions of the CRLM, peri-tumoural and background liver with a limit set at 500 FOVs per tissue.

RNA QC was carried out as per NanoString guidelines using both count matrices and cell sizes. Cells were removed if they breached the following: Cells containing fewer than 50 detected transcripts or displaying abnormally large segmented areas (>40,000 µm²) were flagged and excluded. FOV-level QC involved identifying FOVs with reduced signals or biased gene expression profiles. Cells in these flagged FOVs were excluded from all data objects.

We used Nanostrings in-built Spatial Informatics Portal (SIP) to segment cells prior to downstream analysis. Nuclear and membrane stains are used based on the CellPose algorithm^66^, to perform accurate segmentations. We used the ‘large mixed with small cells’ preset which should be ideal for hepatocytes alongside immune cells, with ML parameters refined based on visual inspection of each FOV.

After quality control, a total of 1,930,656 cells, encompassing 1,845 FOVs, were retained for normalisation and analysis. Samples from different slides were then integrated into a single dataset, and batch effects were corrected using the harmony package (v1.2.3)^67^.

### CosMx normalisation and phenotyping

Target counts were normalised by dividing the total counts of each gene within a cell by the total counts of all genes in that same cell and then multiplying by the average total counts per cell across the entire dataset. UMAP was performed using normalised and square root transformed expression values for all genes. Supervised clustering was carried out using the InSituType package (v2.0) using the reference cell profiles provided by Bruker^68^. The resulting assignments were refined based on marker gene expression. Validation was performed through examination of protein and marker gene expression and the spatial tissue distribution of the cells. Subset-level clustering was performed on pre-filtered expression data containing only the relevant cell types. Further unsupervised clustering was performed using InSituType to identify transcriptionally distinct groups, which were subsequently annotated based on marker gene expression.

### CosMx-specific analyses

#### Region assignment

A Seurat object was constructed using the expression matrix, annotated cell identities, and corresponding metadata using the Seurat package (v5.3.0)^69^. Tumour, peritumour, and healthy regions were defined on a slide-wise basis based on the Euclidean distance of each cell from the nearest cancer cell (Malignant A-B). Cells located within 50 µm of a malignant epithelial cell were designated as the tumour region, those within 50-500 µm as the peritumour region, and >500 µm as background liver. Region assignment was cross-referenced to the original H&E image and fluorescence images.

### Gene signature scores

Gene signature scores were calculated for curated gene sets using a custom function based on Seurat’s ’AddModuleScore’ methodology. For each gene set, control genes were randomly sampled from 24 expression bins based on average expression levels. Per-cell scores were then computed as the average expression of the signature genes minus that of the control genes, averaged across 25 control sets to correct for differences in baseline expression levels. The resulting scores reflect the relative enrichment of each gene signature at the single-cell level. The Hypoxia gene expression score used in figure 6N consisted of the common hypoxia gene set from reference^59^. T-cell exhaustion genesets are referenced in Figure 6.

### Distance-signature score gradients

Signature scores were modelled as a smooth function of the Euclidean distance from the nearest tumour cell. For each slide, as well as across all slides combined, generalised additive models were fitted using the ‘bam’ function from the mgcv package (v1.9.1)^70^. The fitted curves and corresponding 95% confidence intervals were plotted to visualise the continuous variation in signature scores with increasing distance from the tumour region. Analyses were restricted to a common distance range shared among slides to enable direct cross-slide comparison.

### Spatial transcript preprocessing and polygon segmentation

Cell boundaries were reconstructed using CellPoly, which infers polygonal outlines for individual cells based on transcript spatial coordinates to capture cellular morphology and spatial organisation. The resulting segmentation maps were visualised as polygon overlays for representative FOVs.

### Pseudotime Trajectory analysis

Pseudotime analysis was performed using Monocle3, with trajectories rooted in indicated cell types and tissue regions. Pseudotime values were compared across cell types and tissue regions to characterise cell state transitions.

### Cell-cell communication

Cell-cell communication was analysed using the CellChat (v2.2.0)^41^. Separate spatial CellChat objects were generated for tumour, peritumour, and healthy regions. Communication probabilities were calculated using the ‘computeCommunProb’ and ‘computeCommunProbPathway’ function, with an interaction range of 100 µm and a contact range of 10 µm to account for diffusion-mediated and direct cell-cell interactions, respectively.

### scRNA-seq processing and analysis

Two independent scRNA-seq datasets of colorectal cancer liver metastasis and matched normal liver were obtained^71,72^. QC was performed according to the parameters reported in the respective studies removing low-quality cells characterised by low gene counts or high mitochondrial gene expression. Datasets were normalised with the Seurat (v5.3.0) function ’SCTransform’^69^. Processed samples were merged, and batch effects corrected using Harmony (v1.2.3)^67^. The corrected embeddings were used for UMAP visualisation. Unsupervised clustering was performed using ’FindNeighbors’ and ’FindClusters’ to identify distinct cell clusters and cell types annotated based on marker genes.

For downstream analyses, gene expression matrices were log-normalised and scaled using the Seurat functions ’NormalizeData’ and ’ScaleData’. Genes of interest were visualised using violin plots. Curated gene signatures were quantified using the Seurat function ’AddModuleScore’, allowing the assessment of pathway-level activities at the single-cell level.

### Broad spatial analysis methods

#### Neighbourhood Generation

A radius-based QC step filtered spatially isolated cells likely representing segmentation noise or debris. Cells with <3 cells within a fixed Euclidean radius of 100 centroids (65um after scaling) were removed ensuring that retained cells had >2 neighbours. Following Schürch *et al*.^13^, per-cell neighbourhoods were constructed independently in each dataset (Cell DIVE and CosMx). Within each ROI, we used scikit-learn’s NearestNeighbors (v1.6.1)^73^ to find for every cell, its 10 closest neighbours and counted how many of each cell type appeared among those neighbours. This count vector defined the cell’s neighbourhood composition, and stacking these vectors yielded the per-cell neighbourhood feature matrix^13^. Slide-level batch effects were corrected using ComBat (scanpy v1.11.3), while preserving the region structure (covariates=region)^74,75^. Batch-corrected matrices were clustered with K-means, with the number of clusters selected by an elbow analysis of within-cluster inertia across k=1−15. Cluster identities were defined by the z-scored mean frequency of each cell type among member neighbourhoods and used to annotate the per-cell metadata. Relative enrichment and depletion of cell types in each CN compared to their global prevalence across the tissue was imputed. Missing or ill-posed values (NaN or infinite) were replaced with zeros to ensure numerical stability.

### UMAP generation

At the ROI level, we computed the proportion of cells assigned to each neighbourhood cluster, applied ComBat (scanpy) and standardisation (StandardScaler, scikit-learn), and embedded using UMAP (umap-learn v0.5.7; n_neighbors=50, min_dist=0.5, spread=2.0, random_state=42)^76,77^. Two-dimensional embeddings were visualised with ROIs coloured by tissue region and by slide to assess region separation and residual batch structure.

### Cross-Pair Correlation Function (PCF)

We used the Muspan spatial-statistics package (v1.1.1) to compute cross-pair correlation functions (cross-PCF), gij(r), for every ROI in each dataset^78^. ROI-level curves were then pooled within each cohort (Cell DIVE and CosMx), collapsing across all slides and anatomical regions to generate cohort-level summaries. For each ROI, per-cell coordinates (in micrometres) were registered in a muspan domain alongside categorical cell_type labels, and boundaries were estimated with an alpha-shape hull (α=95 μm) with a 10 µm boundary exclusion for edge correction. In a small number of ROIs (n = 3), where the α = 95 μm hull was suboptimal, we re-estimated the boundary with α = 120 μm and recomputed PCFs for those ROIs only. We exhaustively enumerated all ordered cell-type pairs (including self–self) and evaluated cross-PCFs on a common distance grid (0–250 µm; 5 µm steps; 10 µm annuli). Pairs undefined in a given ROI (one or both cell types absent) were written as NaN and omitted from aggregation using NaN-aware means and standard deviations. Outputs from each ROI were recorded under undirected pair labels (“A ⟷ B”), zero-distance artefacts were suppressed (exclude_zero=True), and ROI computations were parallelised per slide. For cohort summaries, curves were aligned to the common grid and averaged only across ROIs in which the pair occurred. Uncertainty was visualised with approximate 95% confidence bands using the normal approximation μ±1.96 σ/√n, where n is the number of contributing ROIs per pair. For display, we generated overlay plots of pre-specified cell-type pairs, showing cohort-level mean gij(r) curves with the 95% confidence bands and the complete randomness reference line at gij(r) = 1. The same workflow was applied independently to the Cell DIVE and CosMx cohorts, yielding whole-cohort cross-PCF summaries that indicate cell-cell clustering (gij(r)>1), randomness (gij(r)=1), and exclusion (gij(r)<1) across spatial scales.

### Topographical Data Analysis (TDA)

TDA was applied to neighbourhood-labelled point clouds in each ROI within the Cell DIVE dataset. Two modes were analysed: single-cluster (centroids of all cells from one neighbourhood cluster) and pairwise-cluster (centroids of all cells from both members of an unordered cluster pair, pooled into a single point set). Vietoris–Rips filtrations were computed with muspan (v1.1.1) using a maximum distance of 707 µm (the diagonal of a 500 µm × 500 µm ROI), and persistent homology was evaluated up to dimension 1 to obtain H0 (connected components) and H1 (loops) barcodes^79^. Analyses were restricted to clusters or cluster pairs with at least three cells per ROI, and persistence diagrams for each ROI were saved for all valid clusters and pairs in that ROI.

TDA persistence diagrams were vectorised into interpretable features^80^, including means, standard deviations and selected percentiles of births, deaths, lifespans and midpoints, plus persistence entropy. To emphasise biologically salient structure, we additionally derived “significant-bar” features after thresholding at ≥50 µm for H0 ≥10 µm for H1. For H0, the significant-bar count was incremented by one to reflect the number of components present just before merges at the 50 µm threshold. ROIs were retained if at least one targeted cluster or cluster pair exhibited a significant H0 component. The final ROI-level panel (1520 ROIs) comprised significant bar counts for H0 and H1, and the means and standard deviations of significant H0/H1 lifespans and significant H1 birth times. UMAPs were generated using the top features ranked by random-forest importance (mean decrease in impurity; Cell DIVE: N = 30). For hypothesis testing, ROI-level feature values were averaged across ROIs from the same slide within the same tissue region to yield one slide-level value per region per slide.

Random forest feature redundancy was screened using pairwise Spearman correlations and Benjamini–Hochberg adjustment; “strong” correlation was predefined as R2≥0.7 (4.7% of tests), supporting the full selected feature panel for classification^81^. Region discrimination was evaluated with random forests trained on ROI-level TDA features (scikit-learn v1.6.1, 1,000 trees) using grouped five-fold cross-validation by slide^82^. Accuracy is reported as mean ± standard deviation across folds, and features were ranked by mean decrease in impurity.

To visualise the spatial pattern of targeted neighbourhood clusters within individual ROIs, we generated ROI-level scatter plots from the per-cell metadata containing cluster labels. Clusters 0, 4, 5, and 7 were highlighted using distinct colours, while all other clusters were rendered light grey to provide context (matplotlib v3.10.1). To visualise H0 structure at the biologically motivated threshold (50 µm), we produced ROI-level component maps for selected neighbourhood clusters. For a chosen ROI and cluster, cells were subset, and a undirected graph was built by connecting points within 50 µm (scikit-learn v1.6.1, radius_neighbors_graph), and connected components were identified (SciPy v1.15.2, connected_components)^83^. Each component was plotted in a distinct colour with all other cells shown in light grey for context, using equal axis scaling in micrometres.

### SCIMAP cell-cell interaction

To quantify short-range spatial interactions independent of tissue region, we pooled all ROIs across the CosMx cohort and computed 10-nearest-neighbour (10-NN) interactions per slide with scimap (v2.3.4)^64^. Per-cell coordinates were loaded into an AnnData object with imageid=slide_ID. We then ran sm.tl.spatial_interaction with method="knn", knn=10, permutation=1000, and pval_method="zscore", yielding slide-level, directional Z-scores and permutation p-values for every ordered pair *i→j* (enrichment if Z>0, avoidance if Z<0).

Directional outputs were exported to long format, then symmetrised within each slide by averaging *i→j* and *j→i* scores (and the corresponding per-slide p-values).

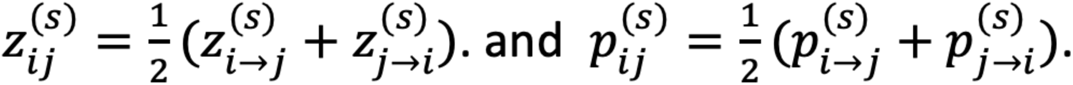

Across slides, the cohort summary for each unordered cell-type pair used the mean symmetrised Z-score, and a conservative aggregate p-value given by the maximum across slides. Pairs were displayed in non-grey colours only if the per-slide (symmetrised) p-value was <0.05 on every slide. Final figures show a lower-triangular heatmap ordered by a curated cell-type list, with a centred diverging colour map (saturating at ±1 Z), and non-significant cells were masked in grey.

### Statistical analysis

Between-region differences for absolute cell and neighbourhood proportions and for selected TDA features were tested at the slide level after averaging across ROIs within each tissue region (CRLM, peri-tumoural liver, background liver) using paired Wilcoxon signed-rank tests with Benjamini–Hochberg false-discovery-rate (FDR < 0.05).

Slide-level means as thin box-and-whisker plots with region-specific colours (interquartile range (IQR) boxes, whiskers to 1.5×IQR, median line shown) (matplotlib v3.10.1). Individual slide values were overlaid as jittered black points to visualise dispersion. Outlying values were labelled by ID when they lay beyond the 1.5×IQR whisker limits.

## Results

### Evaluation of T-cell extravasation in perfused human CRLM, peri-tumoural and background liver

T-lymphocytes are excluded from microsatellite stable (MSS) CRLM, which typically demonstrate an immune cold microenvironment, but the exclusion mechanisms are incompletely understood. We hypothesised that impaired lymphocyte extravasation might contribute to T-cell exclusion. To this end, we developed a disease model system, where surgically resected human hemi-livers containing CRLM are maintained using normothermic perfusion (Fig. 1A). Such models have demonstrated maintenance of the CRLM tumour microenvironment (TME) for up to 40 hours^17^. During a 24-hour period, long enough to compare lymphocyte extravasation into the liver^18^ and stroma-rich tumours^19^, we perfused specimens (supplementary table 1 for clinical details) with oxygenated, nutrient-supplemented venous (portal) and arterial blood under physiological pressures and flows (Fig. 1B), maintaining perfusate biochemistry through hepatic function (Fig. 1C). Within 2 hours of perfusion, we inoculated CFSE labelled lymphocytes from ABO-compatible buffy coats into the perfusion fluid. We were able to detect labelled T-, B- and NK-cells in the perfusate following inoculation using flow cytometry, with the vast majority (>95%) representing CD4 and CD8 T-cells (Supplementary Figure 1). By experimental endpoint, we detected significantly fewer extravasated CFSE^+^ cells in the CRLM compared to the liver (Fig. 1D-E). To confirm that FACS-detected lymphocytes had extravasated and ensure adequate CRLM perfusion, we imaged the fluorescent cells alongside infused Tomato Lectin to label perfused vasculature, comparing perfused regions of background, uninvolved vascular segments with peri-tumoural liver and the CRLM (Fig. 1F-G). Notably, the proportion of extravasated CFSE positive cells was greater within the peri-tumoural liver compared to the CRLM or background liver (Fig. 1G-H), indicating that impaired extravasation contributes to CRLM T-cell exclusion and that circulating lymphocytes primarily extravasate in peri-tumoural liver.

**Figure 1.**
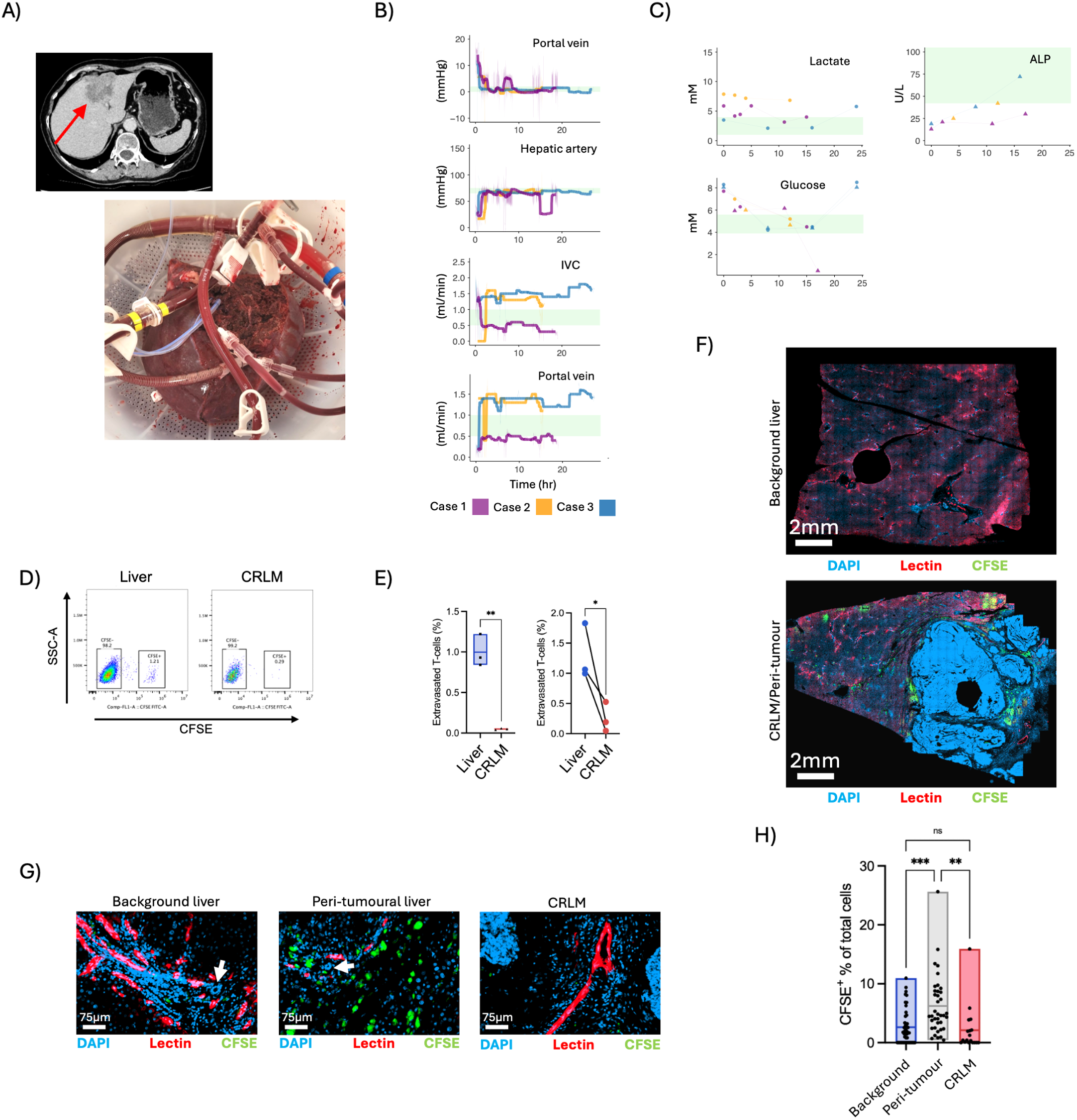
Failure of T-cell extravasation in CRLM. A) Computed tomography image of CRLM (arrow) and resulting explanted left hemi-liver cannulated to enable normothermic perfusion. B-C) Physiological and biochemical readouts from 3 hemi-livers inoculated with CFSE-labelled T-lymphocytes. Green shades represent physiological ranges. D) FACS of disaggregated peri-tumoural liver and CRLM showing detection of extravasated CFSE-labelled T-lymphocytes in the liver, with fewer in the CRLM. E) Quantification of extravasated CFSE^+^ T-lymphocytes in the liver and CRLM across 3 samples from a single perfusion experiment (left) and paired analysis of extravasated T-lymphocyte proportions in liver and CRLM across 3 perfused livers (right). Paired T-test throughout. F) Whole-slide fluorescence image of background liver taken from an uninvolved vascular segment (above) and the CRLM/peri-tumoural liver (below) at the end of the hemi-liver perfusion experiment. Tomato lectin (red), CFSE and DAPI, nuclear label are shown. G) High-magnification images taken from (F) demonstrating examples of portal triads at the indicated locations, identified morphological through the presence of biliary structures (white arrows) and perfused (Tomato Lectin^+^) vascular channels in keeping hepatic arterial and portal venous branches. H) Quantification of extravasated CFSE^+^ T-lymphocyte numbers in the background liver (45 ROIs), peri-tumoural liver (38 ROIs) and CRLM (18 ROIs)(paired T-test). ALP alkaline phosphatase, SSC-A side scatter.

### A spatial transcriptomic CRLM liver atlas

To understand the molecular basis of T-cell exclusion, we generated a single-cell spatial transcriptomic atlas from four replacement CRLM of matched size and chemotherapy regimen (Supplementary Figure 2A, Supplementary Table 2). We segmented 1,930,656 cells (35 cell types)(Fig. 2A, Supplementary Figure 2B-C),) finding conserved proportions across samples (Fig. 2B and Supplementary Figure 2D)^20^. Cells expressed lineage-specific genes (Fig. 2C and Supplementary Figure 3C) and demonstrated appropriate phenotype-specific size parameters (Fig. 2D), supporting our segmentation algorithm. The liver showed cell and transcriptome-level zonation across 4 hepatocyte subtypes (Fig. 2D-E and Supplementary Figure 4A), whilst cholangiocytes expressing hallmark genes *KRT7* and *DEFB1* localised to epithelial structures at the portal triad identified at protein level (Supplementary Figure 4B). The K-nearest neighbour (KNN) method grouped cell phenotypes into biologically relevant neighbourhoods (cN0-cN9, Fig. 2F). CosMx cN2 contained T and B-cells in addition to plasmacytoid and conventional dendritic cells (pDC and cDC) indicating that it is a hub of antigen presentation and adaptive immune response. B-cell and DC presence within cN2 indicates potential antigen presentation and antibody production; features akin to tertiary lymphoid structures (TLS)^21^.

**Figure 2.**
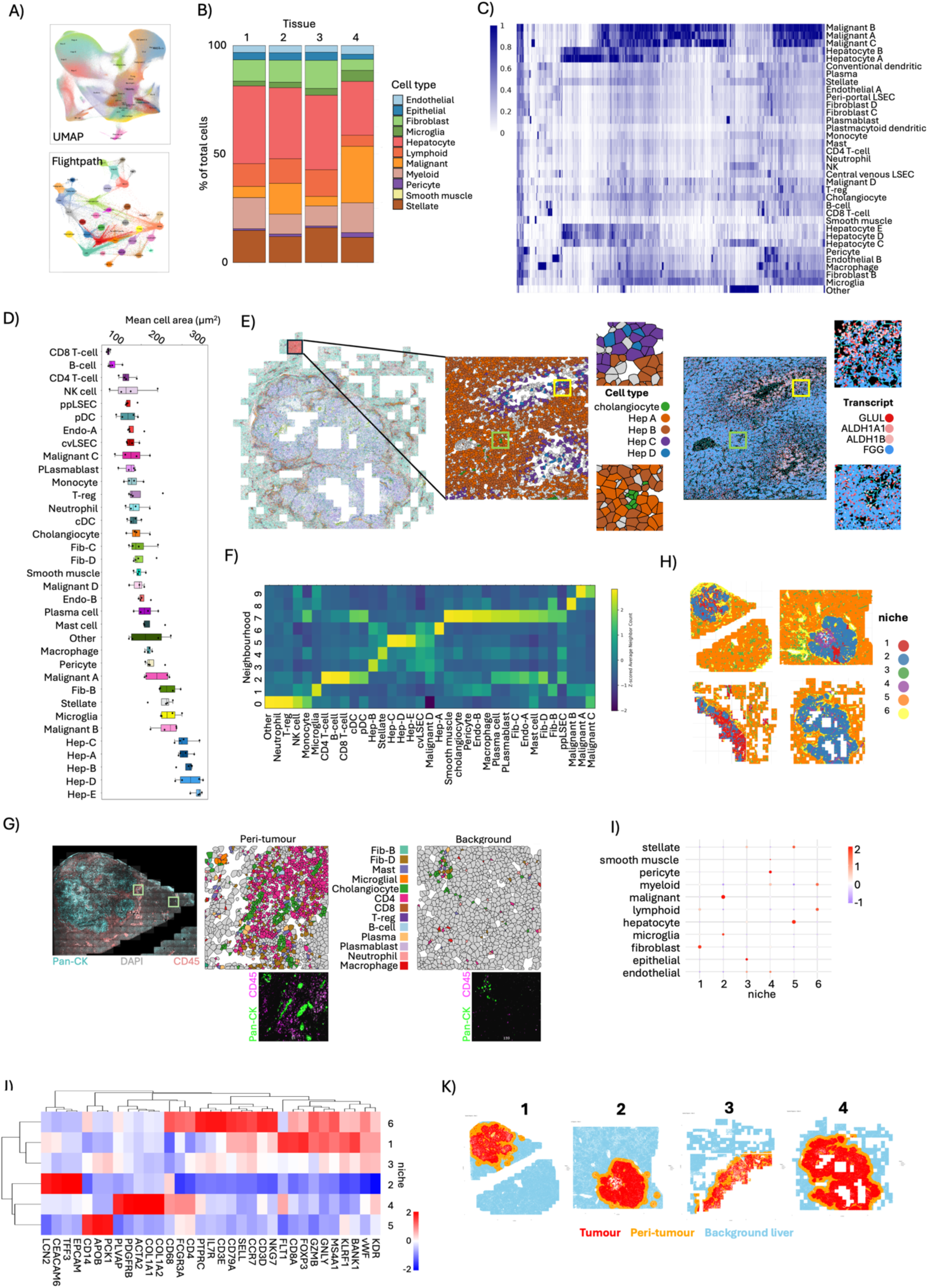
**A spatial transcriptomic CRLM liver atlas**. A) UMAP and flightpath plots demonstrating the results of cell clustering within the spatial transcriptomics (CosMx) dataset (enlarged in Supplementary Figure 3A-B). B) Stacked bar charts of broad cell types in each of the 4 samples showing the proportion of each cell type across the whole specimen. C) Heatmap of gene expression for every gene in the CosMx dataset for each cell type. D) Box-whisker plots of median cell size for each phenotype across the entire CosMx dataset. E) Cell types mapped spatially with magnification of the indicated area demonstrating cell segmentation and colour coding of 4 hepatocyte subsets and cholangiocytes (green) demonstrating zonation with regard hepatocyte distribution. Non hepatocyte/cholangiocyte cell types shaded in grey. The same areas is then shown to the right, but with colour-coded transcript spots representing the locations of the individual RNA molecules demonstrating hepatic zonation with regards transcript distribution. Given the large size of this sample, FOVs of the peritumour and CRLM were prioritised for transcript imaging, as the maximum FOV number suggested by the manufacturer was reached. F) Heatmap showing the 10 most frequently occurring cellular neighbourhoods across all samples determined using the KNN method. G) Whole-slide CosMx immunofluorescence image for the indicated markers (left), with selected CosMx Field of Views (FOVs) from the peri-tumour and background liver demonstrating cell segmentations colour-coded to cell phenotypes. Panel below shows immunofluorescence of the same FOVs demonstrating pan-CK^+^ cholangiocytes whose positions correspond to the cell assignments above. H) Spatial distribution of 6 cell ‘niches’ across the 4 tissues. I) Dot plot of broad cell type frequency for each niche. J) Heatmap demonstrating expression of the indicated genes within each spatial niche. K) Tissue segmentation masks for each sample demonstrating CRLM, peri-tumoural liver and background liver regions.

Although TLS are identified in primary colorectal cancer^22,23^, our spatial segmentation maps did not identify B-cell germinal centres (Fig. 2G), indicating that these are likely lymphoid aggregates rather than mature TLS, supporting that lymphoid aggregates surrounding CRLM rarely represent mature TLS^16^. To investigate how cellular transcriptome changes in different tissue regions, we defined tissue niches from broad cell-type assignments (this is as opposed to the neighbourhood method which considers individual cell type). The resulting niches mapping to their predicted origins in the CRLM or liver (Fig. 2H-I) and bulk transcriptome displayed expression of lineage-specific genes with niche dependency (Fig. 2J). The niches were then combined to generate tumour, peri-tumoural liver and background liver Region of Interest (ROI)s (Fig. 2K). Neighbourhood proportions within different areas of the tissue demonstrated that fibroblast and macrophage neighbourhoods (cN7) are found in both the peri-tumoural liver and CRLM, the lymphoid neighbourhood (cN2) predominantly in the peri-tumoural liver, a neutrophil and Treg-enriched neighbourhood (cN0) in the CRLM, and hepatocyte neighbourhoods (cN3, cN5, cN6) in the peri-tumoural and background liver compartments (Supplementary Figure 2E). Our analysis pipeline thus provides a detailed visualisation of tissue cellular diversity, highlighting the peri-tumoural liver as an area of significant immune infiltration, consistent with the observations in perfused livers where CFSE-labelled lymphocytes extravasated predominantly within the peri-tumoural liver.

### Endothelial anergy in CRLM endothelium

As endothelia regulate T-cell entry into tissues in response to inflammation, we reasoned that differences in endothelial inflammatory signalling in the CRLM and peri-tumoural liver may underlie differences in lymphocyte extravasation and accumulation at these sites. We defined 4 endothelial cell subsets in the CosMx data, including central venous (cv-) and peri-portal (pp-) liver sinusoidal endothelia (LSECs) (Fig. 3A) in the background liver (Fig. 3B-C), endothelial B (Endo-B) predominantly in the CRLM and peri-tumoural liver and Endo-A distributed throughout the tissue regions (Fig. 3C). As well as demonstrating spatial specialisation, endothelial subtypes are defined by specific transcriptional patterns (Supplementary Figure 5A). Pseudotime analysis indicated that Endo-B arises from LSECs, with Endo-A an intermediary state (Fig. 3D and Supplementary Figure 5B), supporting that replacement CRLM co-opt LSEC populations and direct their differentiation to a pro-tumorigenic phenotype as a source of angiogenic vasculature^24,25^. Intra-tumoural endothelia expressed ECM interaction, hypoxia and angiogenic pathways, whilst peri-tumoural endothelia expressed TNF, NFkB, STING, nod-like receptor (NLR) and inflammasome signatures (Fig. 3E) indicating a response to cellular damage. This was apparent when using distance as a continuous variable, where endothelial inflammation peaked in the peri-tumoural region (Fig. 3F). Endothelia therefore shift from a non-inflamed state in the normal liver to a state of inflammation in the peri-tumoural liver and a return to non-inflamed state in the tumour. Toll-like receptor (TLR) signalling remains high in the background liver, in keeping with the LSECs role in pathogen and toxin scavenging^26^.

**Figure 3.**
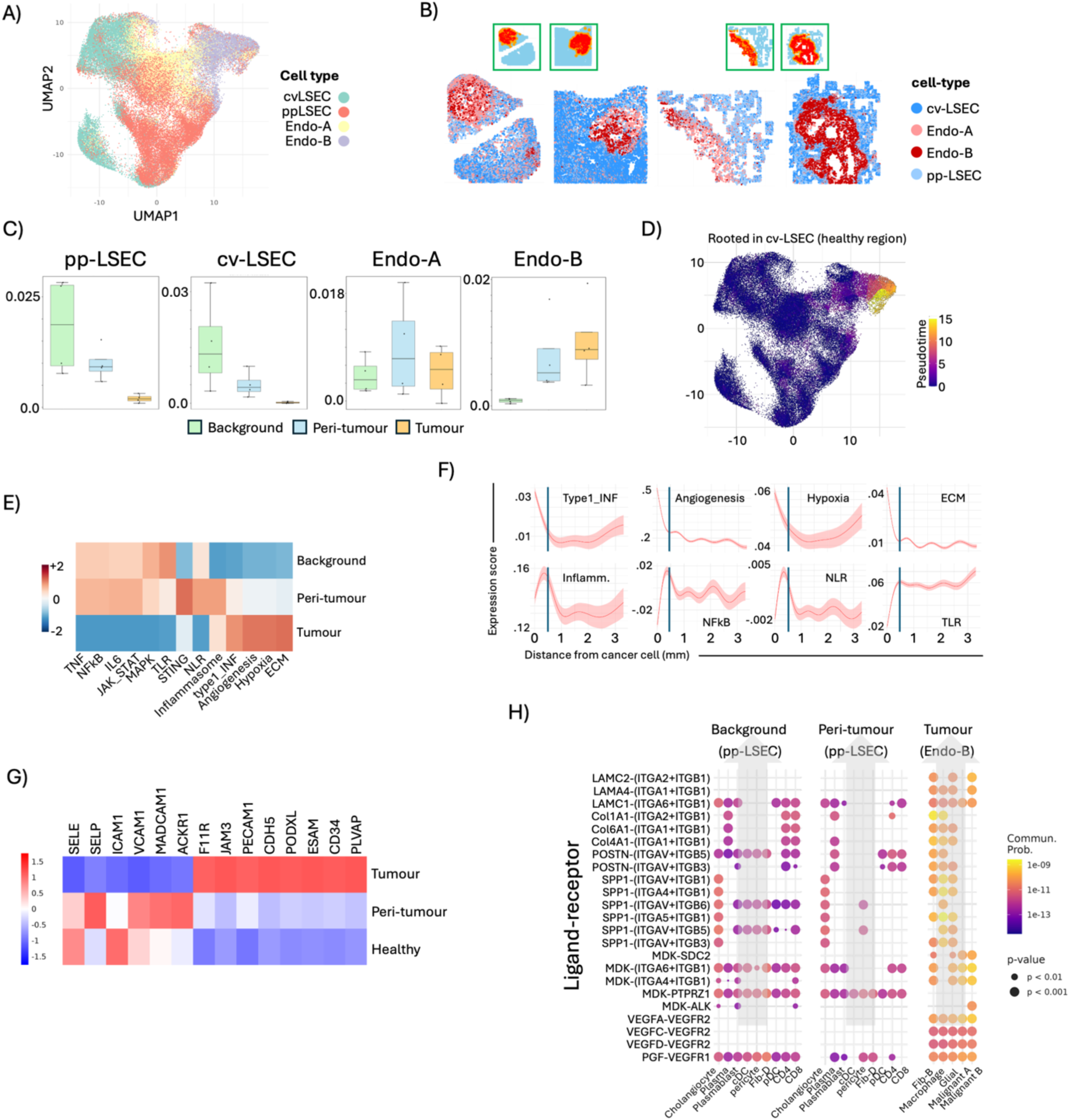
Endothelial anergy in CRLM endothelia. A) UMAP of endothelial subsets. B) Spatial map of the 4 tissues showing the position of the 4 different endothelial cell types. C) Box-whisker of each endothelial subtype across the indicated regions. D) Pseudotime trajectory analysis (Monocle3) of the 4 endothelial populations rooted from the cv-LSEC specifically within the healthy liver. E) Heatmap of the indicated transcriptional signatures of endothelial cells within the different tissue regions. The 4 endothelial cell subsets have been pooled. F) Graphs showing transcriptional signature expression (y-axis) in endothelial cells with distance from the nearest malignant cell (x-axis) in mm. The blue vertical line indicates the distance boundary between the peri-tumour and background liver (i.e. 500µm). Dark curves represent mean and shaded regions around the curve represent the confidence interval. Gene expression score is averaged for each cell across all tissues. G) Heatmap of expression of selective adhesion molecule genes in all endothelial cells within the indicated tissue regions. H) Bubble plot of ligand-receptor interactions with different endothelial subtypes within specific tissue locations as the receiver cell (top), ligand-receptor pairs on the y-axis and emitting cell types on the x-axis. Grey arrows show the direction of signalling. Colour and size of dots represent communication strength (relative probability) and p-value is calculated using a random permutation test.

Inflammation drives adhesion molecule expression in endothelia, and T-cell adhesion molecules *SELP*, *VCAM1*, *MADCAM1* and *ACKR1* were specifically downregulated in CRLM endothelia relative to those in the peri-tumoural liver (Fig. 3G). Conversely, expression of endothelial cell-cell interaction and barrier maintenance genes *JAM1* (*F11R*), *PECAM1*, *CDH5*, *PODXL*, and *PLVAP* were highest in CRLM endothelia (Fig. 3G). By combining 2 publicly available scRNAseq datasets (E-MTAB-12022 and dbGAP identifier phs001818.v3.p1), we also identified higher adhesion molecule expression and upregulation of damage response pathways were demonstrated in background liver endothelia compared to those in the CRLM (Supplementary Figure 5C-E). Ligand-receptor analysis of endothelia segregated to tissue region demonstrated that fibroblast-B (Fib-B), macrophages and cancer cells communicate with Endo-B through ECM basement membrane laminins and collagens (*COL4* and *COL6*), whilst driving angiogenesis through *VEGF, PGF, MDK* and *SPP1* ligands (Fig. 3H), a feature lacking in endothelia within the background and peri-tumoural liver. Lack of a basement membrane is a recognised feature of the hepatic LSEC population, and our findings indicate progressive anergy and capillarisation of the endothelial tree from the peri-tumoural liver to the CRLM, with potential endothelial-ECM interactions likely to provide a further barrier to effective T-cell extravasation^27^. Transcriptional anergy demonstrated here provides a molecular explanation for the failed T-cell extravasation observed in the perfusion platform (Fig. 1). This is the first time endothelial anergy has been identified as a potential mechanism of CRLM T-cell exclusion; a feature that warrants further mechanistic validation to determine whether reversal of endothelial anergy can promote intra-tumoural T-cell extravasation.

### Fibroblast phenotypes in CRLM and peri-tumoural liver

CosMx neighbourhood analysis indicated maintenance of a fibroblast-rich neighbourhood (cN7) from the peri-tumoural liver to the CRLM (Fig. 2F and Supplementary Figure 2E). We identified 3 fibroblast populations (Fib. B-D), with Fib-C and Fib-D demonstrating transcriptional similarity (Fig. 4A). Fib-D and Fib-B displayed strong outgoing signalling capacity in the peri-tumoural liver and CRLM respectively, with neither contributing in the background liver, indicating a spatial component to fibroblast specialisation and that fibroblasts are key regulators of both the CRLM and peri-tumoural microenvironments (Fig. 4B). Fib-B expressed neutrophil chemo-attractants *CXCL1* and *CXCL8*, ECM remodelling genes *TNC* and *LOXL2* and transcription factors *SOX9* and *HMGA1*, which promote activation and fibrosis in cancer and chronic inflammatory states^28,29,30,31^. Fib-C expressed collagen and elastin fibril and ECM modifying genes *Col14a1, LUM, Col6a1, Col6a2, Col6a3* and *DCN* and Fib-D expressed genes involved in TLS formation including *CCL21*^32^, *CCL19*^16^(Fig. 4C). Fib-D also expressed HLA Class II transcripts, indicating capability to present antigen to CD4 T-cells; a feature identified in cancer-associated fibroblasts at other sites^33,34^.

**Figure 4.**
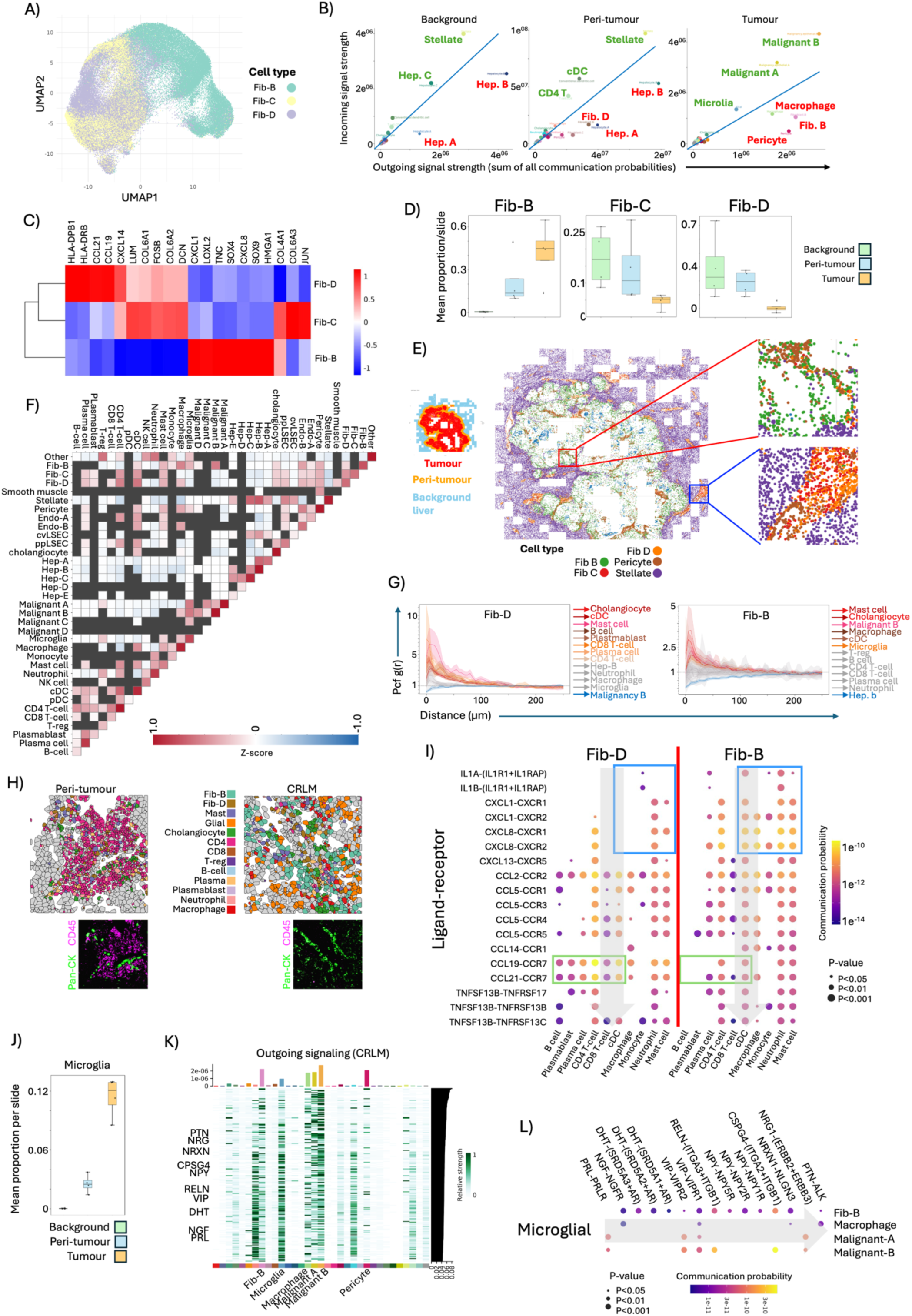
Fibroblast plasticity in CRLM and background liver. A) UMAP of fibroblasts divided into 3 subtypes across the CosMx dataset. B) Incoming and outgoing signalling strengths based on cumulative ligand/receptor expression separated based on tissue region and with strongest incoming/outgoing signalling cell types highlighted. Axes indicate the summation of all incoming or outgoing communication probabilities^41^. Cell types in red are net outgoing signallers and those in green are net signalling receivers. The blue line indicates perfect balance in outgoing and incoming communication probability. Cell types in the bottom left of each graph play lesser roles in incoming/outgoing signalling. C) Heatmap of selected genes from the 3 fibroblast populations. D) Box-whisker plots of the different fibroblast subtypes in the indicated tissue regions. Given small sample size (4 per condition), statistical difference not formally tested. E) Whole tissue spatial map of stromal cells with tissue segmentations shown (left) for reference. Higher magnification (right) demonstrates tumour stroma (top) and portal triad in background liver (bottom). F) SCIMAP single cell-cell spatial proximity plot showing the likelihood of the indicated cell pairs residing within the ten nearest neighbours of each other. Red indicates spatially correlated and blue represents spatial exclusion. G) Pair correlation function (PCF) plots describing the ratio of the observed density of cell pairs g(r) (y-axis), at distance r (x-axis) to the expected density under complete spatial randomness. Displaying Fib-D (left) and Fib-B (right) as the index cell. Curves are coloured red (statistically significant spatial association) grey (no association – random distribution) and blues (statistically significant spatial exclusion). Lines indicate means with 95% CI shaded. CI excluding 1 up to at least 100µm is considered significant. H) Example cell segmentation masks taken from portal triad identified within the peritumoral liver (left) and CRLM (right) regions. Cholangiocytes are identified in each plot, but with a significantly altered immune-stromal infiltrate inclusive of microglial cells, Fib-B and macrophages in the latter. I) Ligand-receptor bubble plot of interactions with Fib-D (left) or Fib-B (right) as the emitter of ligands and various cell types as the receivers (x-axis). Colour and size of dots represent communication strength (relative probability) and p-value is calculated using a random permutation test. Plots limited to the peritumour and CRLM for Fib-D and Fib-B respectively. J) Box-whisker plots showing the proportion of microglial cells in the CRLM, peri-tumour and background liver. K) Plot showing the relative strength of outgoing transcriptional programs from cell types in the CRLM with signalling factors specific to microglial cells indicated on the y-axis. L) Bubble plot showing interactions between microglia as the signalling cell to various receiving cell types (right hand side), for specific ligand-receptor pairs (x-axis, top).

Fib-B represented the greatest fibroblast proportion in CRLM, with Fib-C and Fib-D in the peri-tumoural and background liver, but rarely in CRLM (Fig. 4D-E) indicating that Fib-D phenotypic loss accompanies and may be mechanistically linked to failure of lymphoid aggregate maintenance in CRLM (Fig. 6, below). Other stromal cells (e.g. stellate) were confined to background liver, with pericytes in both CRLM and liver (Fig. 4E). Fib-D were spatially correlated with pp-LSECs and cholangiocytes indicating residence within the portal triad (Fig. 4F-G), but also with plasma cells, cDC’s, CD4 and CD8 T-cells (Fig. 4F), and interaction over longer distances with B-cells (Fig. 4G). Conversely, Fib-B spatially correlated with cancer cells, microglia, monocytes and macrophages, and were spatially inversely correlated with both T-reg and CD4 T-cells (Fig. 4F). Both Fib-B and Fib-D showed strong spatial association with cholangiocytes (Fig. 4G) which were present in the peri-tumoural liver and the CRLM (Fig. 4H). Replacement CRLM thus co-opt and replace the surrounding liver, leaving the biliary tree intact, but altering its immune composition from one of immune activation and exhaustion to myeloid and microglial infiltration (Fig. 4H).

Fib-D expressed key mediators of TLS formation to neighbourhood cells including *CCL19, CCL21* and *CXCL13* (Fig. 4I), whilst Fib-B expressed macrophage recruiting factors *CXCL1, CXCL8, CCL2, CCL5* and IL1 (Fig. 4I). Fib-D communicated extensively with B- and Plasma cells, with Fib-B displaying a complete lack of such communication (Fig. 4I). This indicates that the CRLM becomes devoid of a fibroblast population potentially capable of supporting adaptive anti-tumour immunity and antibody production; key features of functional TLS at cancer sites outside of the liver^21,35^.

Finally, we investigated cells that might drive fibroblast plasticity in the CRLM. We focused on microglia, as they were spatially associated with Fib-B (Fig. 4F-H), located almost exclusively in CRLM (Fig. 4J) and displayed outgoing signalling potential in the CRLM (Fig. 4B). Microglia are the most abundant immune cell type in the central nervous system, sharing transcriptional properties and immunomodulatory functions with macrophages^36^. More recently, they have been identified in the peripheral nervous system^37,38^, although they have not been studies in the context of malignancy outside of CNS cancers. Microglia expressed ligands not identified in other cells spatially linked to Fib-B, but characteristic of microglial cell biology, including nerve growth factor, neuropeptide, dihydrotestosterone and pleiotrophin (Fig. 4K). These factors all drive fibroblast proliferation and collagen production^39,40^, and each interacts with receptors expressed on Fib-B (Fig. 4L), indicating the presence of a microglial-fibroblast signalling axis within the CRLM with the potential to drive fibroblast activation and ECM production. Given no prior examination of microglial cell phenotype in CRLM, these findings need to be validated at protein level across larger patient cohorts before mechanistic studies to investigate whether targeting the microglial-fibroblast signalling axis has the potential for anti-cancer immunomodulation.

### Protein-level spatial validation of CRLM T-cell exclusion

To spatially validate the immune exclusion identified in our perfused liver experiments, we profiled 13 MSS whole-slide CRLM and background liver samples (Fig. 5A and Supplementary Figure 6A), simultaneously visualising 19 proteins (Fig. 5B-C) and phenotyping >10^7^ cells (23 cell types)(Fig. 5D and Supplementary Figure 6B-C). See supplementary table 3 for clinical details. T-cells appeared excluded from CRLM (Supplementary Figure 6D) and resided in peri-tumoural aggregates at the portal triad (Supplementary Figure 7A-B), corroborating the CosMx data. The lymphoid aggregates contained exhausted and, to a lesser extent, proliferating T-cells (Supplementary Figure 8A-B) and portal triad T-cell density decreased with triad distance from the CRLM-liver border indicating a tumour-specific immunological response (Supplementary Figure 8C-E).

**Figure 5.**
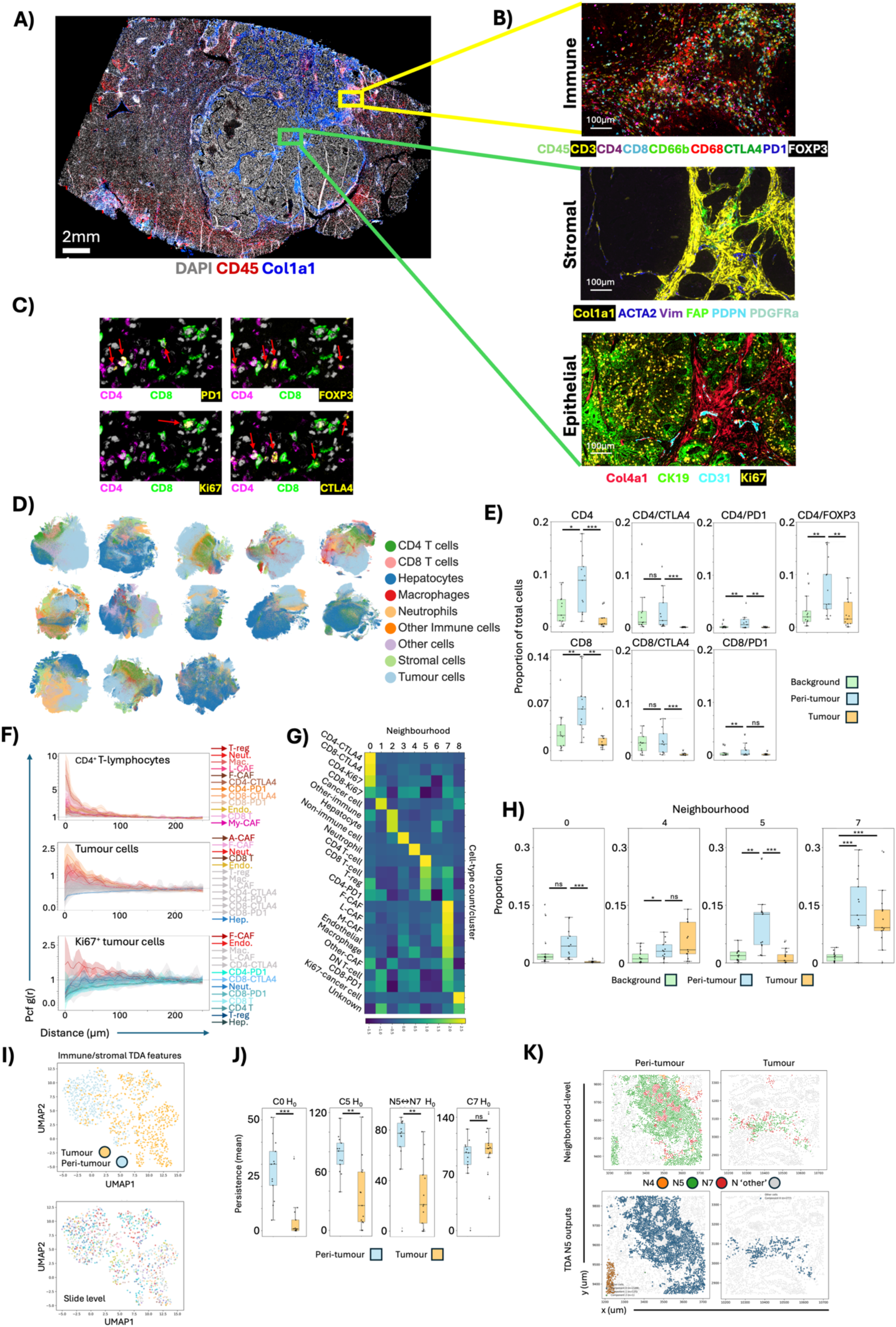
Organised cellular exhaustion niches in the peri-tumoural liver. A) Whole-slide multiplexed immunofluorescence of a CRLM and background liver imaged using the Cell DIVE platform (Leica) with inset (B) demonstrating antibody panels at higher magnification. C) Identification of early exhausted (CTLA4^+^), late exhausted (PD1^+^), proliferating (Ki67^+^) CD4 and CD8 T-cell populations and CD4^+^/FOXP3^+^ T-regs. Red arrows indicate double positive cells. D) Uniform manifold approximation and projection (UMAP)s of broad cell phenotypes for each sample. E) Box-whisker plots of CD4 and CD8 T-cell subsets in the CRLM, peri-tumoural liver (liver tissue ≤500µm of the CRLM edge) and background liver (liver tissue >500µm from the CRLM edge). Individual sample medians from multiple ROIs represented by points, central line represents median, and error bars interquartile range (IQR). Analysed using paired Wilcoxon signed-rank tests at the slide level adjusted for false discovery (Hochberg FDR)^44^. F) PCF plots with settings as per Figure 3G, but using CD4^+^ T-lymphocytes (top), non-proliferating cancer cells (middle) and proliferating cancer cells (bottom) as the index cells. G) Heatmap showing the 9 most frequently occurring cellular neighbourhoods across all samples determined using the KNN method. H) Box-whisker plots of N0, N4, N5 and N7 frequency in the indicated tissue regions across all samples. Presentation and analysis as per (E). I) UMAP of N0, N5 and N7 TDA features selected using Random Forest classifiers applied to vectorised TDA curves describing the structural arrangement of these neighbourhoods in the tumour (orange) and peri-tumoural liver (blue). Lower pane demonstrates the slide that each point from the UMAP was derived, with the 13 slides colour coded demonstrating lack of batch effect and supporting that the tissue region distinguishing TDA features are common between samples. J) Box-whisker plots of the indicated TDA feature persistence in the CRLM and peri-tumoural liver. Presentation and analysis as per (E). K) Spatial map of the peri-tumour (left) and CRLM (right) colour coded by neighbourhood (top) and N5 TDA component (bottom) highlighting breakup of T-cell enriched structures within the CRLM.

We segmented tissues into ‘CRLM’ (bounded by CK19^+^ cells), ‘peri-tumoural’ liver (cells ≤500µm from the CRLM) and ‘background liver’ (cells >500µm from the CRLM) akin to the CosMx analysis. In keeping with our observations (Supplementary Figure 6-8), CD4, and to a lesser extent, CD8 T-cells including early (CTLA4^+^) and terminally (PD1^+^) exhausted and proliferating phenotypes, were most prevalent in the peri-tumoural liver, indicating it is the primary site of immune activation and exhaustion (Fig. 5E)^42,43^ and confirming that lymphocytes are effectively excluded from the CRLM. CD4 T-cells demonstrated significant spatial interaction with other immune cell types across cell-cell signalling distances (<100µm), corroborating the CosMx data (Fig. 5F). Cancer cells, conversely, demonstrated spatial association with fibroblasts and endothelia, with proliferating cancer cells demonstrating strong exclusion of all T-cell subsets (Fig. 5F). Proliferating cancer cells therefore reside in protected, immune-excluded niches and lymphocytes form organised communities in the peri-tumoural liver that are excluded from the CRLM.

K-Nearest Neighbour (KNN) analysis identified cellular neighbourhoods of early exhausted (CTLA4^+^) and proliferating CD4 T-cells (Neighbourhood 0, N0), and of CD4 T-cells, CD8 T-cells, terminally exhausted (PD1^+^) CD4 T-cells and T-regs (N5)(Fig. 5G). N0 and N5 exist in the peri-tumoural liver and are almost completely absent from the CRLM (Fig. 5H). Fibroblast, endothelia, macrophage, double-negative T-cell and terminally exhausted CD8 T-cell clusters (N7) appear at equal proportion in peri-tumoural liver and CRLM (Fig. 5H). Broadly, this validates the spatial transcriptomic observations.

We applied topological data analysis (TDA) to N0, N5 and N7, allowing us to map structural changes in the immune and stromal environments of the CRLM and peri-tumoural liver. TDA distinguishes the frequency of disconnected ‘islands’ and their spatial longevity prior to merging with other islands (H_0_). This structural persistence identifies features of potential biological importance. Random forest of vectorised TDA curves defined the peri-tumour and CRLM with high cross-validated accuracy (72.7±7.4 %)(Fig. 5I), so the CRLM and peri-tumoural liver are identified by the unique structural biology of their stromal populations. The top 30 TDA features were dominated by H_0_ descriptors from N0, N5, and stromal/myeloid rich interactions (e.g. N5↔N7)(Supplementary Figure 9A-B), with each significantly elevated in the peri-tumour (Fig. 5J). Conversely, N7 (fibroblast/macrophage) persistence was equivalent in the CRLM and peri-tumour (Fig. 5J). The CRLM thus maintains myeloid and stromal structures, whilst disrupting the exhausted lymphoid aggregates as it grows into and co-opts peri-tumoural liver (Fig. 5K). These findings identify structural fragmentation of exhausted lymphoid aggregates as the CRLM grows out into the peri-tumoural liver and phenocopies the CosMx data, where fibroblast-cholangiocyte spatial relationships were maintained across the peri-tumoural liver and CRLM, with loss of lymphoid populations and their structural integrity in the latter.

### CD4 T-cell dysfunction in the CRLM and peri-tumoural liver

Our Cell DIVE data identified CD4 T-cell predominance in the peri-tumoural liver, supporting published findings^42,43^. CD4 T-cells are of increasingly recognised importance in the CRLM TME^42^, and display diverse functional repertoires^45,46,47^. In adoptive T-cell trials, neoantigen-specific CD4 T-cell, but not CD8 T-cell proportions correlate with clinical response, pointing to their role in mediating anti-tumour immunity^48^. Because CD4 T-cells mediate hepatic inflammation^49^, maintain TLS through interaction with immune and stromal cells^50^, and were the predominant immune population in the peri-tumoural liver, we decided to return to the spatial transcriptome dataset to understand CD4 subset spatial biology.

Unsupervised clustering of CD4^+^ cells defined 6 new sub-populations (Fig. 6A) including follicular helper (CD4-Tfh) expressing *BCL6, PDCD1* and *CXCR5*, cytotoxic (CD4-Ctl) expressing *GZMH* and *NKG7*, central memory (CD4-Tcm) expressing *CCR7, KLF2* and *IL7R,* T-regulatory (CD4-Treg) expressing *FOXP3, CTLA4* and *TNFRSF4*, stress response state (CD4-Tstr), expressing *HSPA1A, HSPA1B*, and *HSPB1* and naïve CD4 T-cells (CD4-Tn)(Fig. 6B). Pseudotime analysis indicated that CD4-Tstr is a terminal subset (Fig. 6C). CD4-Tcm were found at higher number in the peri-tumoural liver (Fig. 6D-E) and were excluded from cancer cells (Fig. 6F), whereas the CD4-Tstr represented the majority of intra-tumoural CD4 T-cells (Fig. 6D and Supplementary Figure 10A-B) and were the only CD4 subtype spatially correlated with cancer cells (Fig. 6F). Despite variation in the proportion of several CD4 subsets between samples, CD4-Tcm loss and CD4-Tstr gain from the peri-tumour to the CRLM was a common feature across samples (Supplementary Figure 10B). CD4 T-cells displaying a stress response are identified in lung, breast and renal cancers^46^, but not previously in CRLM.

**Figure 6.**
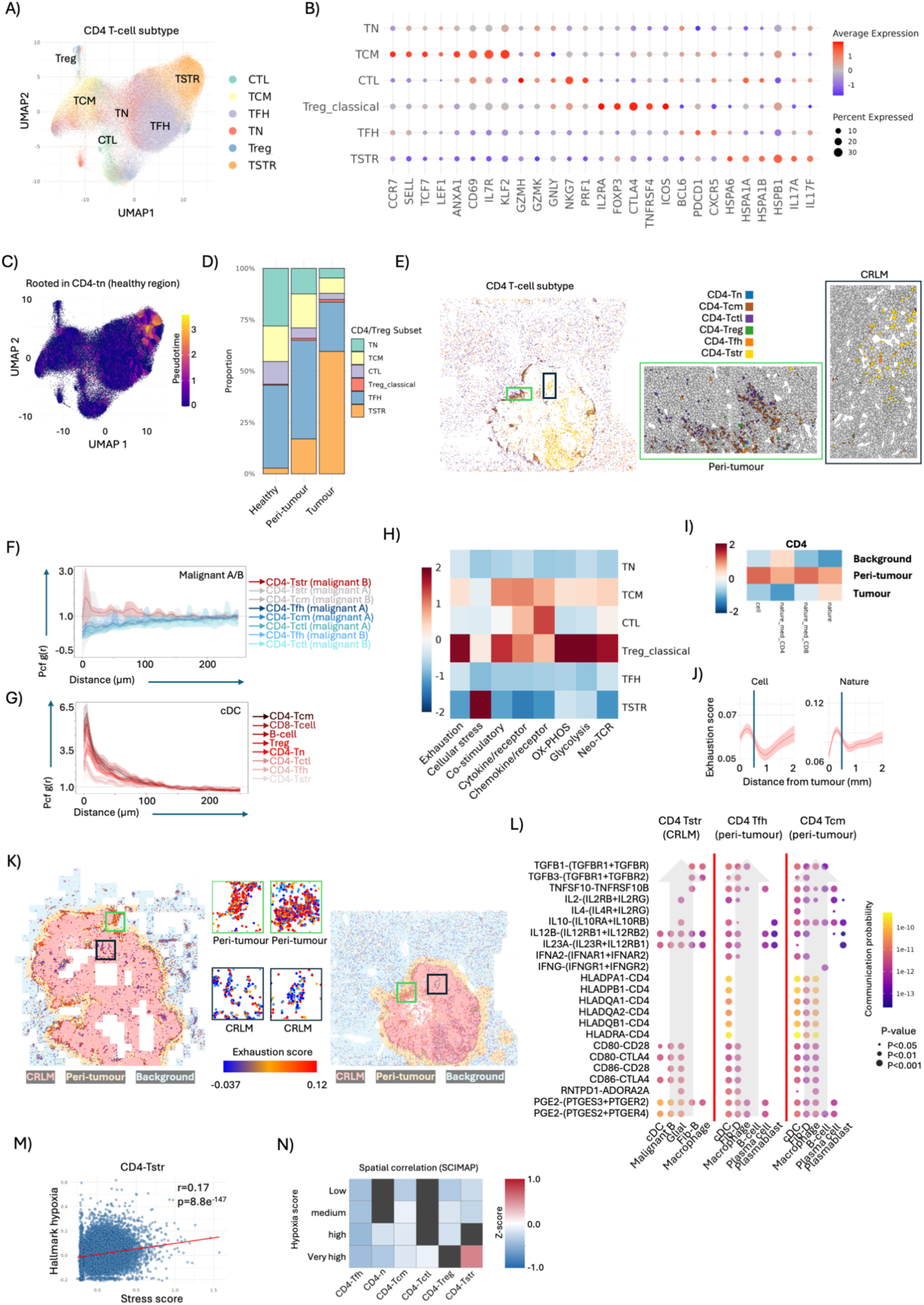
CD4 T-cell dysfunction in the CRLM and peri-tumoural liver. A) UMAP of CD4 T-cell subsets. B) Dot plot of marker expression displaying hallmark genes for each CD4 T-cell subset. C) Pseudotime trajectory analysis of CD4 T-cell subsets. D) Stacked bar plot of the proportion of different CD4 T-cells subsets in the indicated tissue regions. E) Spatial map with CD4 T-cell subsets colour-coded see Figure 2(K) for CRLM/peritumour/background boundaries. F-G) PCF plots showing the spatial relationships between CD4 T-cell subsets and malignant A/B cells (F) and cDC (G) with settings as per Figure 4G. H) Heatmap of the expression of different transcriptional pathways (x-axis) in each of the CD4 T-cell subsets. I) Heatmap of the indicated T-cell exhaustion signatures (x-axis) in CD4 T-cells from the three tissue regions. Exhaustion signatures were from references ^56^(Cell), ^57^(Nature Medicine) and ^58^(Nature). J) Graphs showing exhaustion signature expression in CD4 T-cells with distance from the nearest malignant cell (x-axis) in mm. Settings as per Figure 3F. The signature source is indicated above each graph. K) Spatial map of CD4 T-cells in 2 of the samples colour-coded based on enrichment of the Cell exhaustion signature score from (I). The CRLM, peri-tumour and background liver are overlayed in pink, yellow and blue respectively. Inset high-magnification boxes show heavily exhausted cells in the peri-tumoural liver (top boxes) and lower levels of exhaustion in the CRLM (lower boxes). L) Bubble plot of ligand-receptor interactions with settings as per Figure 3H, showing signalling from different cell types (x-axis) converging on the indicated CD4 T-cell subsets (top). Grey arrows show the direction of signalling. M) Scatter plot of transcriptional stress (x-axis) and hypoxia (y-axis) signatures in the CD4-Tstr population. N) SCIMAP single cell-cell spatial proximity plot showing the likelihood of the different CD4 T-cell subsets residing within the ten nearest neighbours of any cell type from the CosMx dataset based on the cells hypoxic state (expression level of hypoxic gene signature from reference^59^). Red indicates spatially correlated and blue represents spatial exclusion.

Central memory T-cells are those that respond to previously encountered antigen. In keeping with this, CD4-Tcm showed the strongest spatial correlation with antigen presenting cells such as cDC (Fig. 6G), and upregulation of a neo-TCR transcriptional pathway indicative of response to neoantigen (Fig. 6H)^51^. CD4-Tcm also displayed cytokine expression and upregulation of co-stimulatory molecules but simultaneously displayed high levels of exhaustion (Fig. 6H). The CD4-Tstr subset lacked cytokine expression capability and demonstrated a low TCR score indicating failure to maintain neoantigen reactive CD4 T-cells within the tumour core either through loss of TCR-reactive clones, or through their reversion to a non-TCR reactive subtype. CD4-Tstr did not display high levels of exhaustion and overall, CD4 T-cell exhaustion levels peaked in the peri-tumoural liver (Fig. 6I-K), where CD4-Tcm and CD4-Tfh are found at high levels. This validates the Cell-DIVE findings and indicates that exhaustion is not a mechanism through which the CRLM drives immune evasion within the tumour core, but that peri-tumoural exhaustion may be brought about through chronic antigen exposure, with CD4 T-cells becoming dysfunctional through alternative pathways in the CRLM, resulting in a cellular stress response.

Supporting these ideas, we found that antigen presenting cells including cDCs, Fib-D and macrophages interact with CD4-Tcm in the peri-tumorual liver through MHC and co-stimulatory molecules *CD80/CD-86* (Fig. 6L). In the peri-tumoural liver CD4-Tcm receive strong incoming signalling through *IL10*, *TGFb1*, *TGFb3* and *TRAIL* (*TNFSF10*), all of which drive exhaustion and (in the case of *TRAIL*), promote T-cell apoptosis^52^. Conversely, CD4-Tstr receive no MHC signalling and are exposed to prostaglandin (*PGE2*) and IL23A by several different cell types. *PGE2* does not affect T-cell priming, but specifically prevents the expansion of effector T-cell populations within the TME^53^ and is a prominent driver of T-cell stress responses^54^. Furthermore, *PGE2* and *IL23A* combine to drive *IL17* expression by CD4 T-cells^55^, a cytokine with both anti- and pro-tumourigenic effects expressed by the CD4-Tstr population (Fig. 6B). Hyoxia is also an important driver of the cellular stress response and a prominent feature of the TME. Notably CD4-Tstr cellular stress response was positively correlated with hypoxia signalling on a single cell level (Fig. 6M), and CD4-Tstr cells but not other CD4 populations demonstrated spatial correlation with the most hypoxic cells across all tissue regions (Fig. 6N) indicating that the CD4-Tstr specifically reside within hypoxic microenvironments.

## Discussion

Here, we demonstrate for the first time, failed T-cell extravasation in human CRLM coupled with endothelial anergy, providing a novel mechanism through which CRLM successfully exclude circulating T-cells. Simultaneously, our data indicate that lymphocytes signposted for extravasation in the peri-tumoural liver through endothelial inflammation and fibroblast-derived chemokine expression, become exhausted before reaching the CRLM. This process likely provides a conducive soil for CRLM outgrowth, which it accomplishes through co-option of the hepatic sinusoid; a process we conclusively demonstrate through identification of retained biliary epithelia within the CRLM indicative of portal triad structural maintenance. Their identification enabled us to show that the outgrowing CRLM replace exhausted peri-portal lymphoid aggregates with macrophages recruited predominantly through the secretory activity of plastic fibroblast populations driven by microglia. CD4 T-cells retained from the peri-tumoural liver, or migrating into established CRLM reside within hypoxic niches and display a stress response, impaired T-cell receptor and cytokine activity. Future work is needed to dissect the kinetics through which T-cell populations become exhausted or stressed such that therapeutics to reverse these processes can be developed.

In summary, coupling of a novel *in vivo* human tumour perfusion system with a large single-cell spatial transcriptomic dataset from whole-slide specimens validated through mIF has provided a wealth of findings of relevance for the therapeutic targeting of immune evasion in CRLM. Limiting our analysis to microsatellite stable, replacement-type CRLM treated with neoadjuvant chemotherapy enabled us to study the most common clinical scenario and helped to reduce confounding in view of the small sample size; the major limitation of our work. As such, profiling a larger cohort will be essential to confirm the presence of the identified cell states across patient subsets and to correlate their presence and spatial transcriptional state with clinical outcome. Nonetheless, our findings represent the most detailed spatial analysis of the CRLM microenvironment and background liver to date, providing a roadmap for future mechanistic studies aiming to sensitise to immunotherapeutic approaches such as adoptive T-cell therapies and ICB.

## The work was funded by

CRUK Discovery Grant DRCPGM-Jun25/100008

The Lee Placito Fellowship

Theolytics Ltd funded components of the organ perfusion experiments

The Oxford Cancer Centre

## Author Contribution

Wan, Cheng, Wu, Mian: Data analysis/interpretation, data acquisition

Johnson, Bottomley: Data analysis/interpretation, manuscript writing

Lee, Hazini, Moradi, Friesen, Caviezel, Abbas, Sadik, Lakha, K. Jones, Kumaran, R. Jones, Coles: Data acquisition

Bashford-Rogers: Writing, supervision/funding

Gordon-Weeks, Fisher, Carlisle, Coussios, Seymour: Conceptualisation/design, manuscript writing, data acquisition, supervision/funding

## Supplementary Figures

**Supplementary Figure 1.**
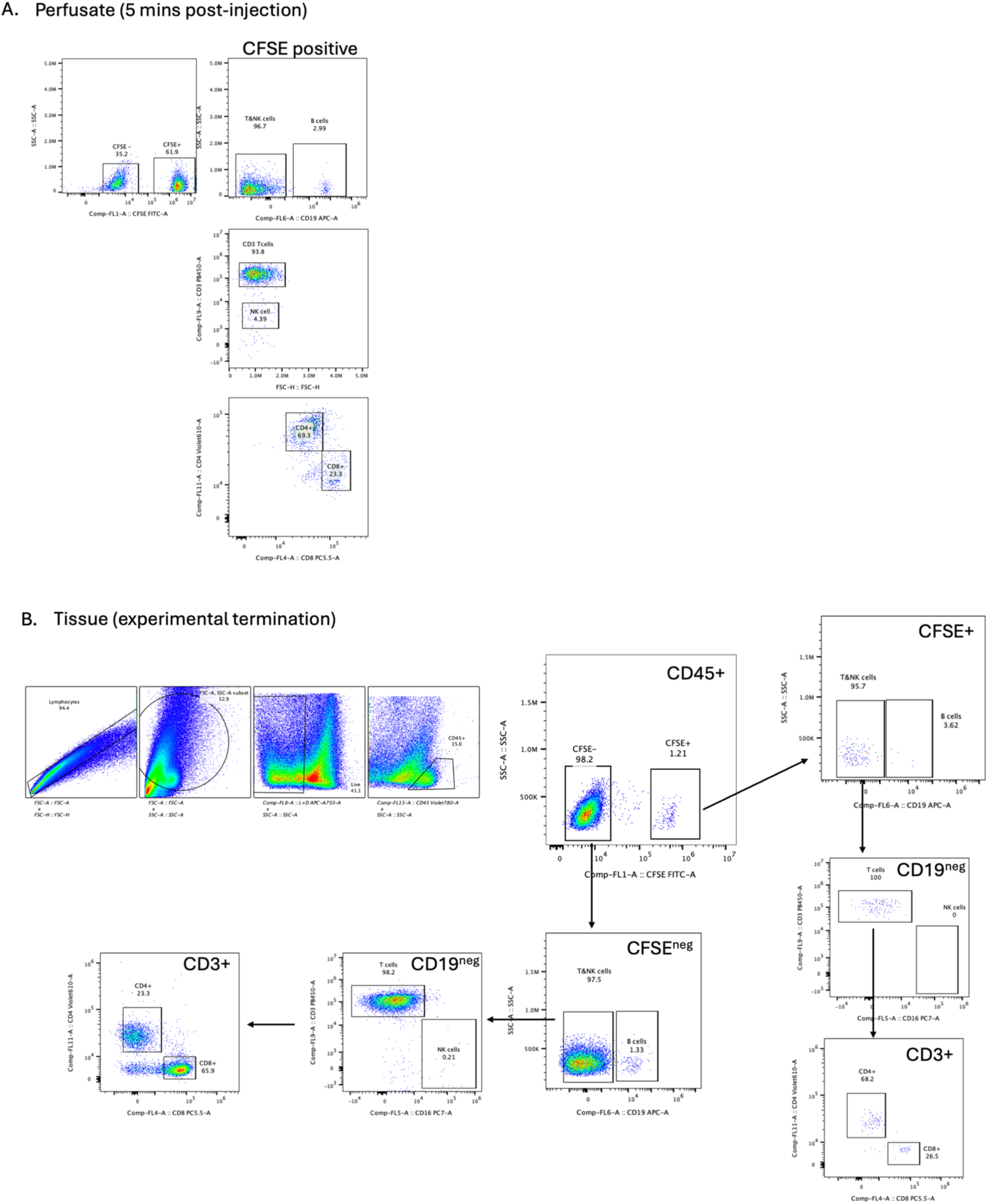
Gating strategy for flow cytometry. **A)** FACS plots of perfusion samples post red cell lysis demonstrating expression of CFSE in a proportion of the circulating immune compartment. Subsequent plots are pre-gated on the CFSE positive population and demonstrate presence of B-cell, CD4/CD8 T-cell and NK cell markers. The perfusate sample was taken 5 minutes following lymphocyte inoculation. **B)** FACS plots of disaggregated tissue samples taken at the experimental endpoint demonstrating gating strategy and showing that the CFSE expressing population that extravasates within the liver consists of lymphocytes expressing CD3, CD4 and CD8 with a lesser population expressing CD19.

**Supplementary Figure 2.**
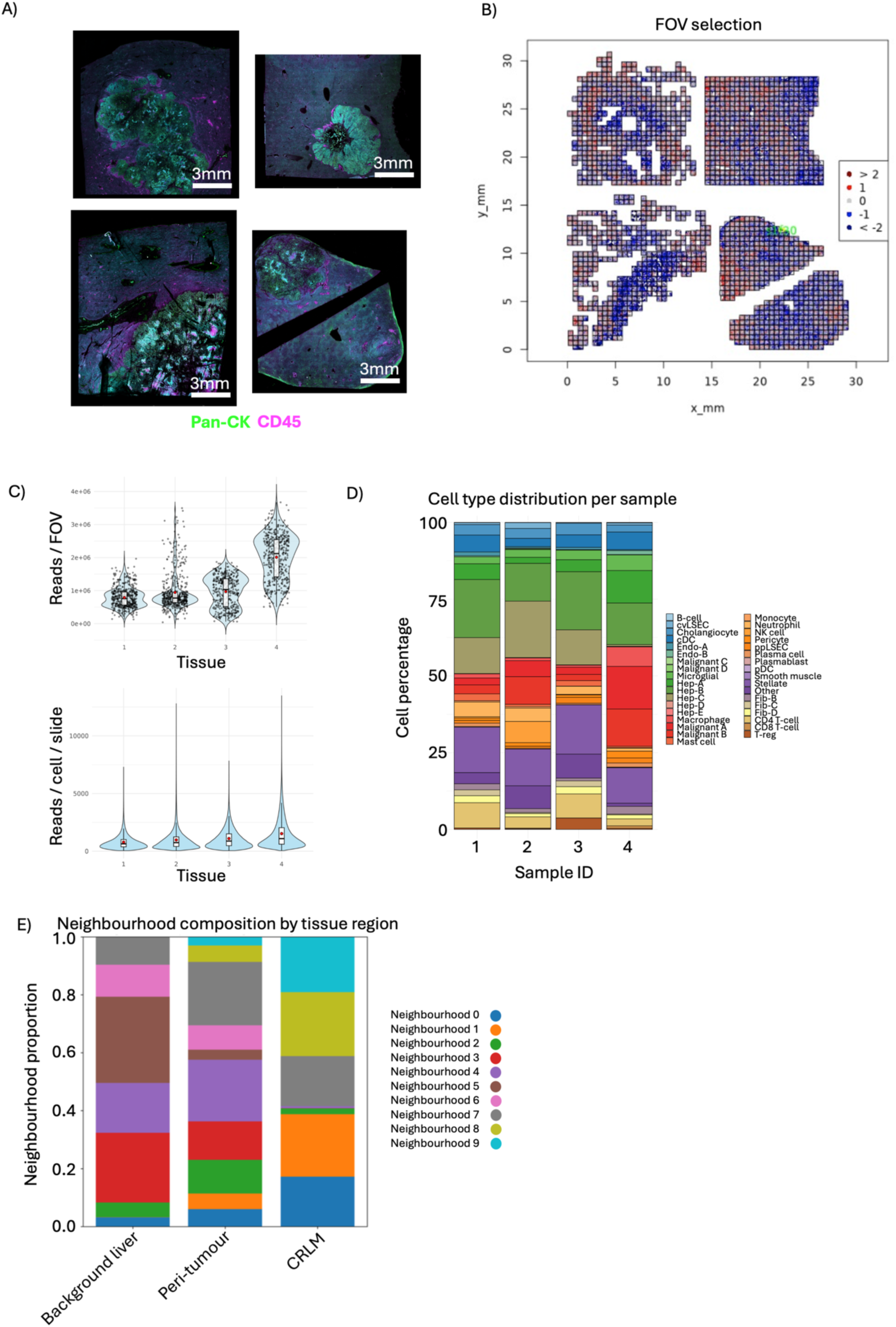
CosMx platform QC. **A)** Immunofluorescence for indicated markers from CosMx platform on the 4 tissues used to generate the spatial transcriptomic atlas. **B)** Example of a flagged FOV exhibiting diminished total counts or biased gene expression profiles leading to exclusion. **C)** Violin plots demonstrating the number of reads per FOV for each tissue (top) and the number of reads per cell (bottom) across the entire CosMx dataset. **D)** Stacked bargraphs showing the proportion of each cell type across the 4 specimens within the CosMx dataset. Red dots represent corresponding group mean. **E)** Stacked bargraphs showing the proportion of each neighbourhood defined within the CosMx dataset (Figure 2F) across the three tissue regions.

**Supplementary Figure 3.**
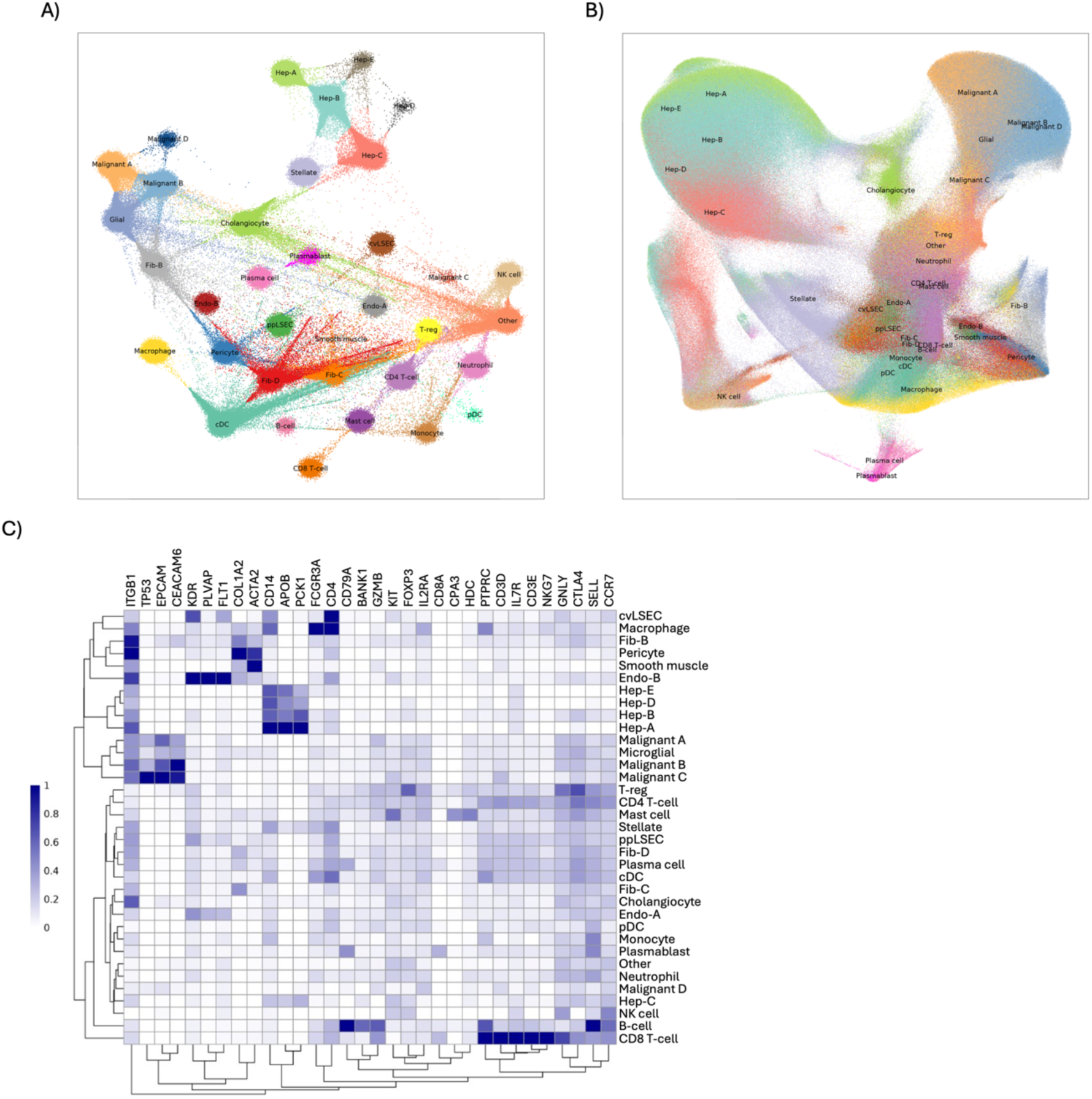
Gene expression characteristics of single cell transcriptomic atlas. **A)** Flightpath plot enlarged image from Figure 2(A). **B)** UMAP cell clustering of CosMx data enlarged from Figure 2(A). **C)** Heatmap of selected hallmark genes defining each cell type in the CosMx dataset.

**Supplementary Figure 4.**
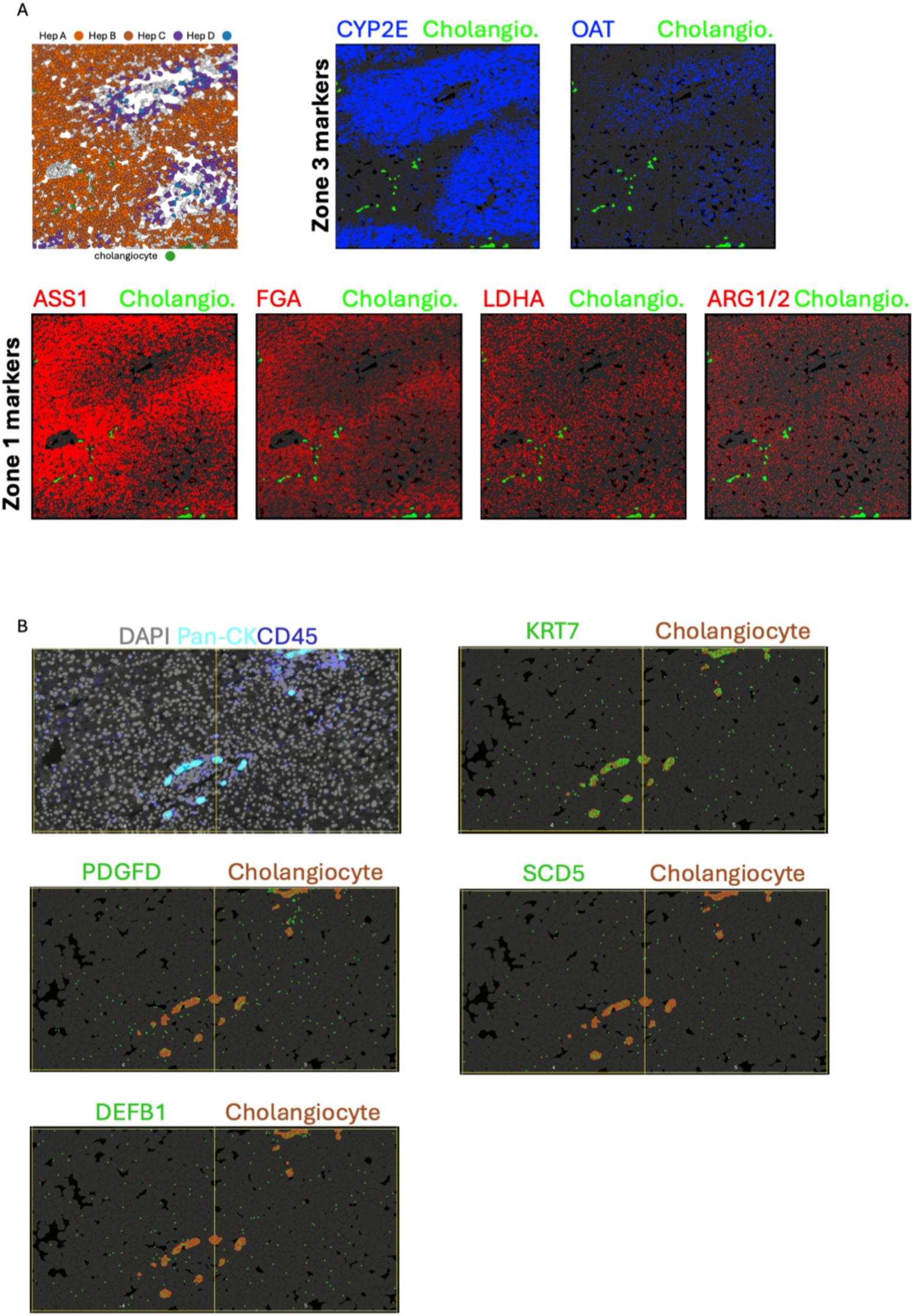
Validation of phenotyping strategy in hepatocytes and cholangiocytes. **A)** Spatial cell phenotyping from CosMx data with hepatocyte subsets colour-coded based on clustering from figure 2A. Hepatocytes A and B are predominantly in a peri-portal location (sinusoidal zone 1), whilst hepatocytes C and D are found nearer the central vein (sinusoidal zone 3). Cholangiocytes demonstrated in green. Subsequent panels demonstrate CosMx-detected transcript positions for genes known to regulate peri-portal (red) or pericentral (blue) functions superimposed on cholangiocyte segmentations (green). These data spatially validate the phenotypic assignment of hepatocyte subsets. **B)** CosMx immunofluorescence for the indicated markers showing positive pan-cytokeratin staining in cell clusters within the background liver. Subsequent panels show CosMx phenotyped cholangiocytes (brown) alongside detections of the canonical cholangiocyte markers KRT7, PDGFD, SCD5 and DEFB1, all of which are enriched in the cholangiocyte segmentations.

**Supplementary Figure 5.**
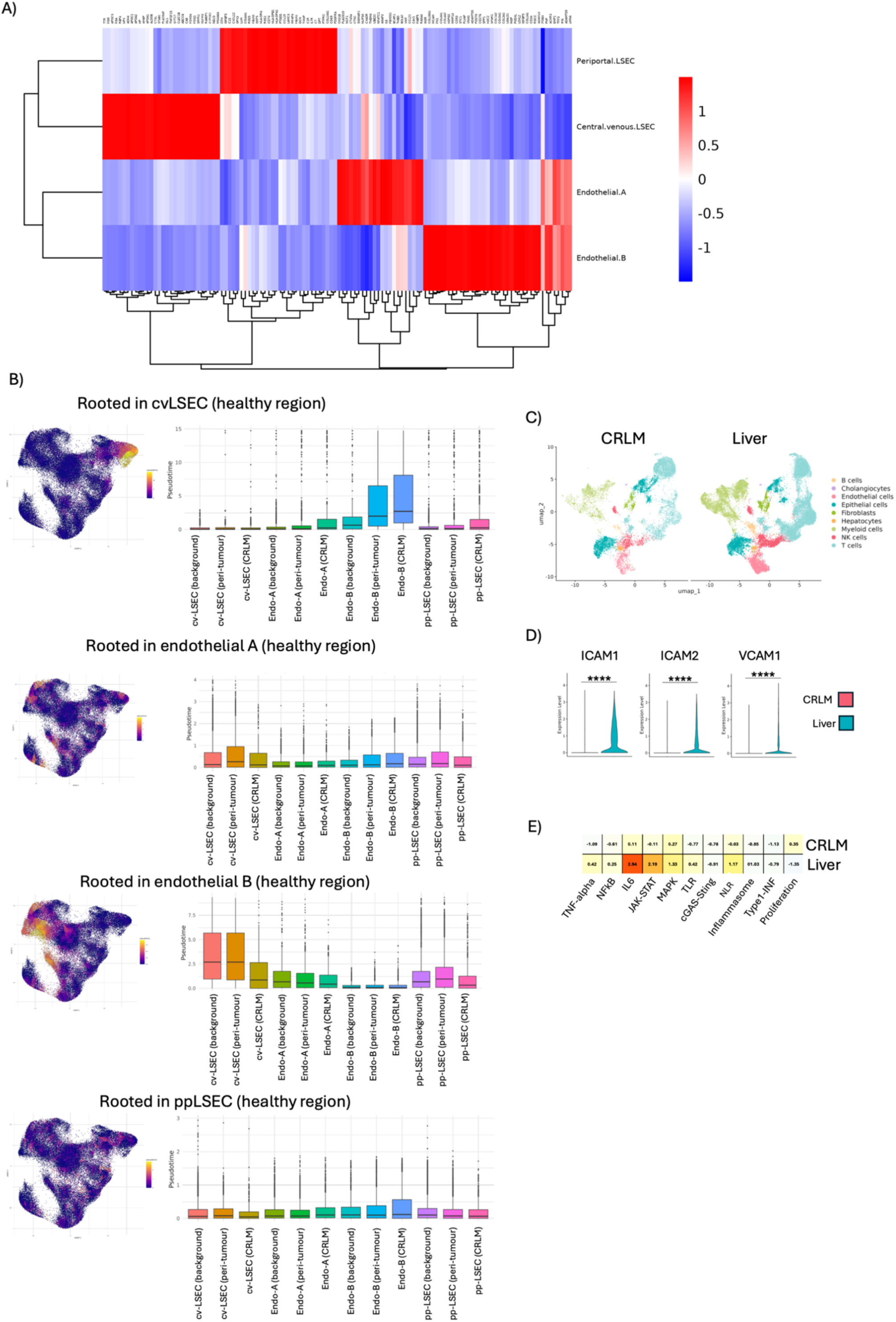
Single cell sequencing of pooled CRLM and background liver validates CRLM endothelial anergy and reduced adhesion molecule expression. **A)** Heatmap of top 30 differentially expressed genes in each endothelial cell subtype. **B)** pseudotime plots for endothelial cells rooted in the indicated endothelial subtype and spatial location. **C)** UMAP clustering of CRLM and background liver samples from two batch corrected single cell sequencing datasets with cell types colour coded. **D)** Violin plots of endothelial adhesion molecule expression for the indicated genes in the CRLM and background liver. **E)** Heatmap of the indicated gene expression signatures in endothelial cells from the CRLM and background liver using the same signatures as those in Figure 3.

**Supplementary Figure 6.**
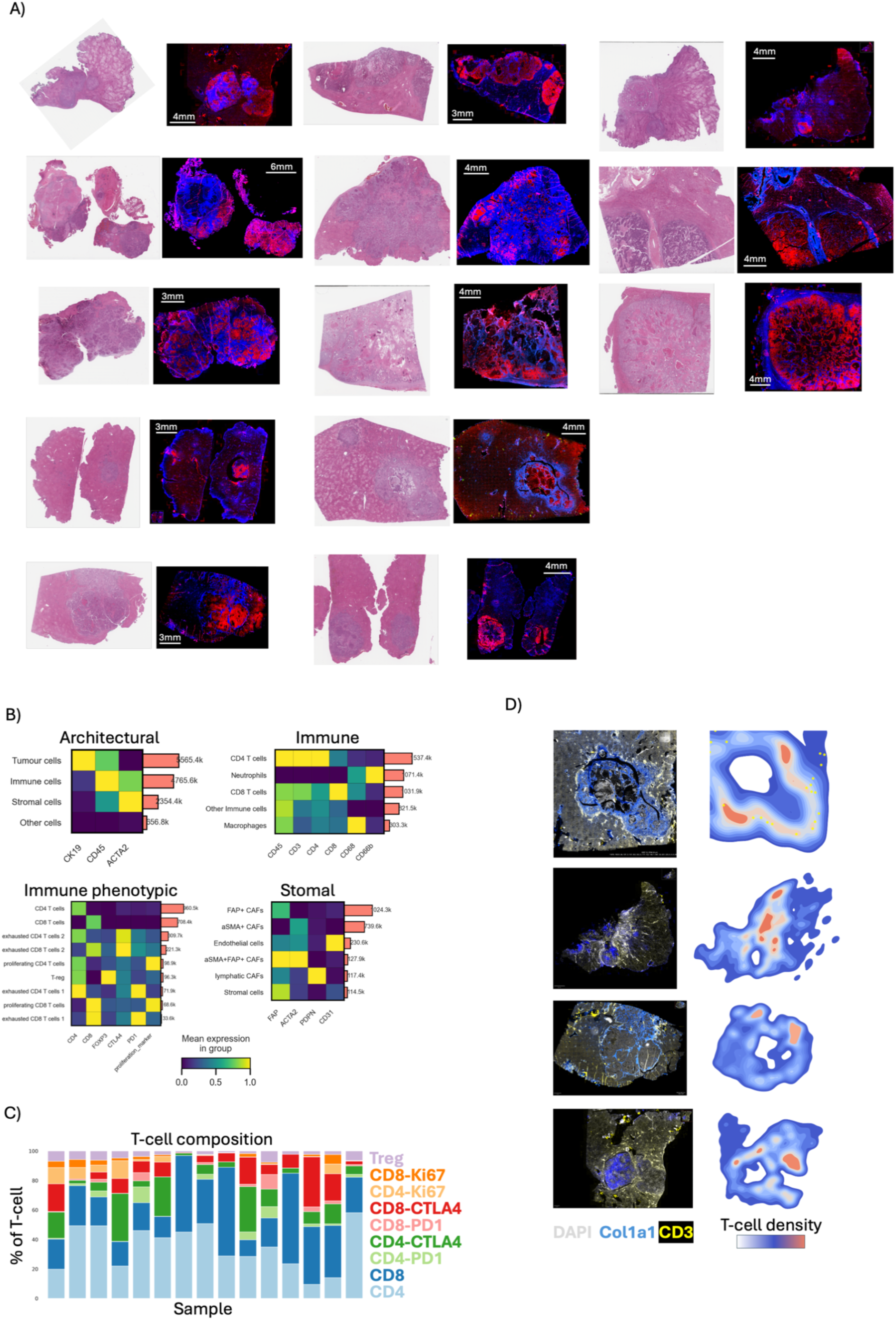
mIF dataset and cell phenotyping strategy. **A)** H&E and matched mIF of the Cell-DIVE dataset stained for CK19 (red) and collagen 1 (blue). **B)** Heatmaps of protein expression level used for phenotyping strategy demonstrating protein marker on the x-axis and cell assignment on the y-axis with numbers of each cell type demonstrated by the pink bars. **C)** Stacked bar graphs of the proportion of different T-cell subsets for each whole slide. **D)** mIF for the indicated markers and match T-cell spatial heatmaps demonstrating exclusion of T-cells from the tumour (indicated by Col1a1 staining).

**Supplementary Figure 7.**
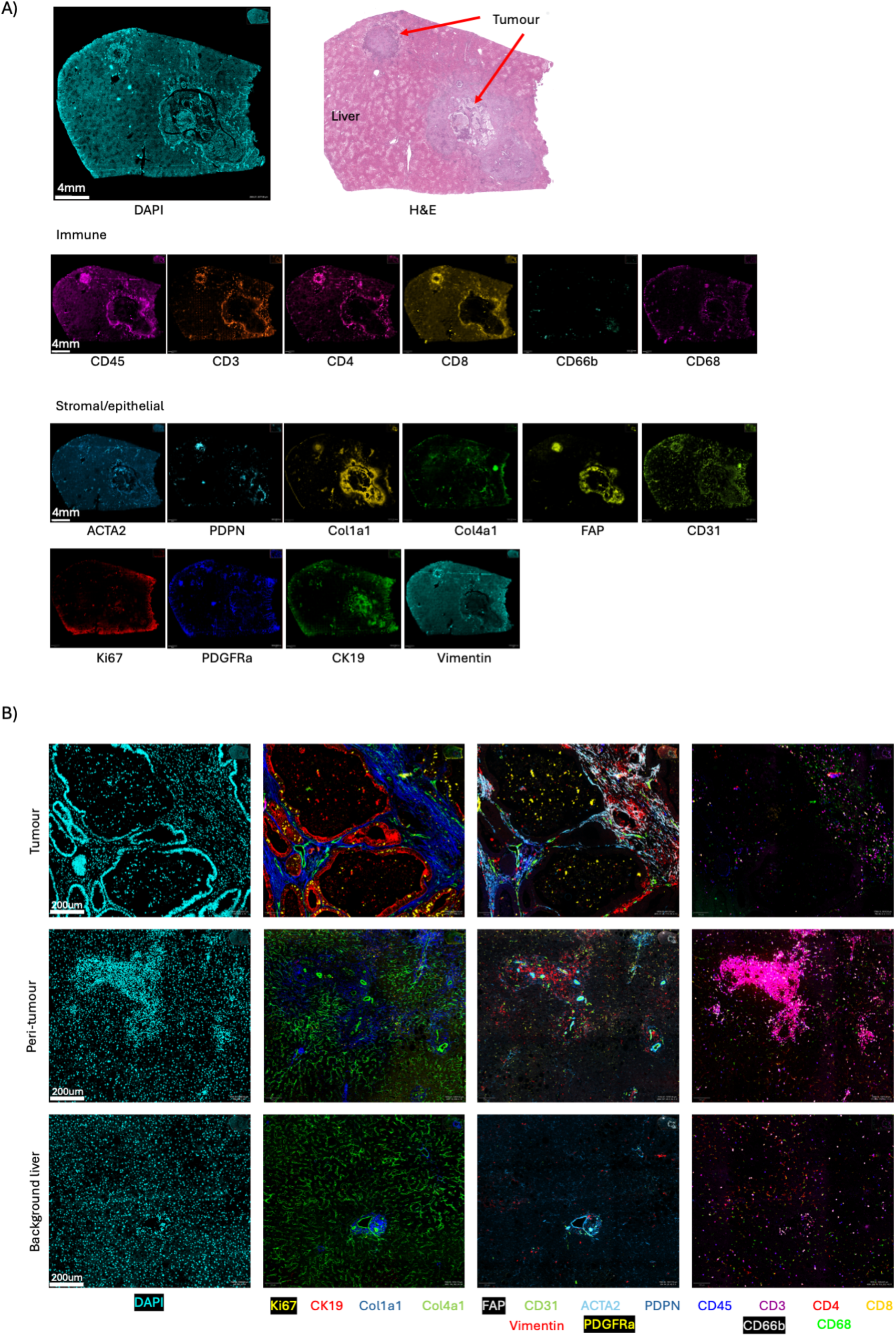
Whole slide marker expression and spatial patterning in the CRLM, peri-tumoural liver and background liver. **A)** Whole slide image of CRLM (tumour) and background liver showing DAPI and matched H&E with tumour positions marked. The distribution of each marker is demonstrated below. **B)** Multiplex for the indicated markers in the CRLM, peri-tumoural liver and background liver demonstrating the organised immune infiltrate in the peri-tumoural liver, which surrounds the portal triad as demonstrated by the presence of CK19^+^ epithelia surrounded by Col4a1 and Col1a1 (peri-tumour, second panel from the left). The CRLM displays dense ECM deposition and proliferation within the malignant epithelial compartment (Ki67^+^). In both the background liver and CRLM, the organised, lymphocytic infiltrate seen in the peri-tumoural liver is lost.

**Supplementary Figure 8.**
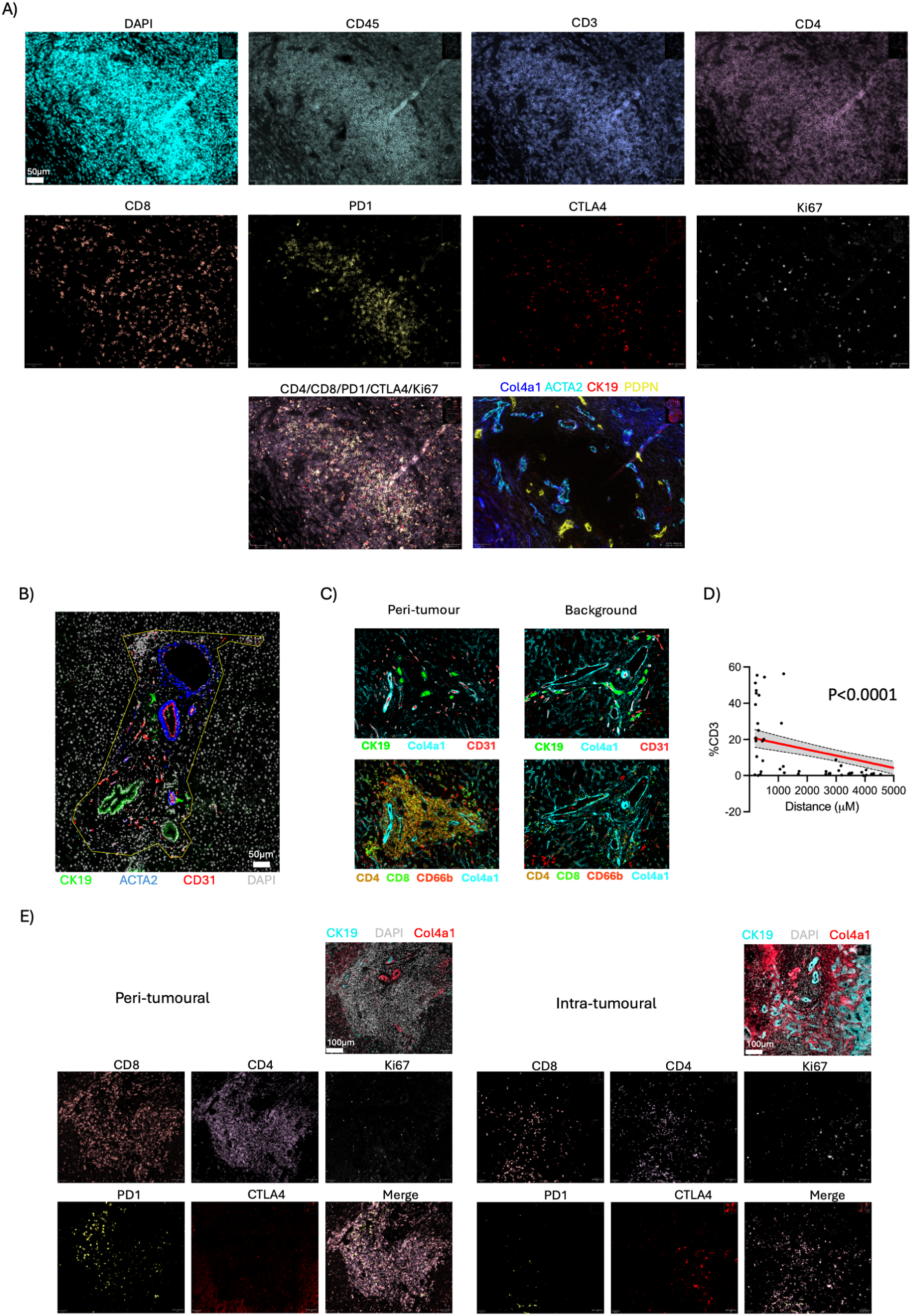
Organised lymphoid aggregates are a feature of the peri-tumoural liver and contain exhausted lymphocytes. **A)** Multiplexed image of lymphoid aggregate in the peri-tumoural liver demonstrating the predominance of CD4 T-cells over CD8. This is a portal triad, as evidenced by spatially matched multiplex demonstrating the presence of CK19^+^ biliary epithelium surrounded by Col4a1 and ACTA2^+^ fibroblasts. **B)** Multiplexed image demonstrating ROI generation used to define portal triad. Biliary tree (CK19), portal vein (CD31) and hepatic artery (CD31 surrounded by ACTA2^+^ myofibroblasts) branches are identified. Resultant ROI demonstrated by yellow border. **C)** High magnification comparison of the portal triad in the peri-tumoural and background liver demonstrating the CD4-rich lymphoid aggregate in the former. **D)** Graph showing correlation between CD3 count (proportion of total segmented cells) per portal triad, y-axis and distance of portal triad from the liver-tumour interface in microns (x-axis). P-value reported for Pearsons correlation coefficient. **E)** Multiplexed image of peri-tumoural liver and CRLM microenvironment for the indicated markers demonstrating organisation of lymphocytes and expression of PD1 in the former.

**Supplementary Figure 9.**
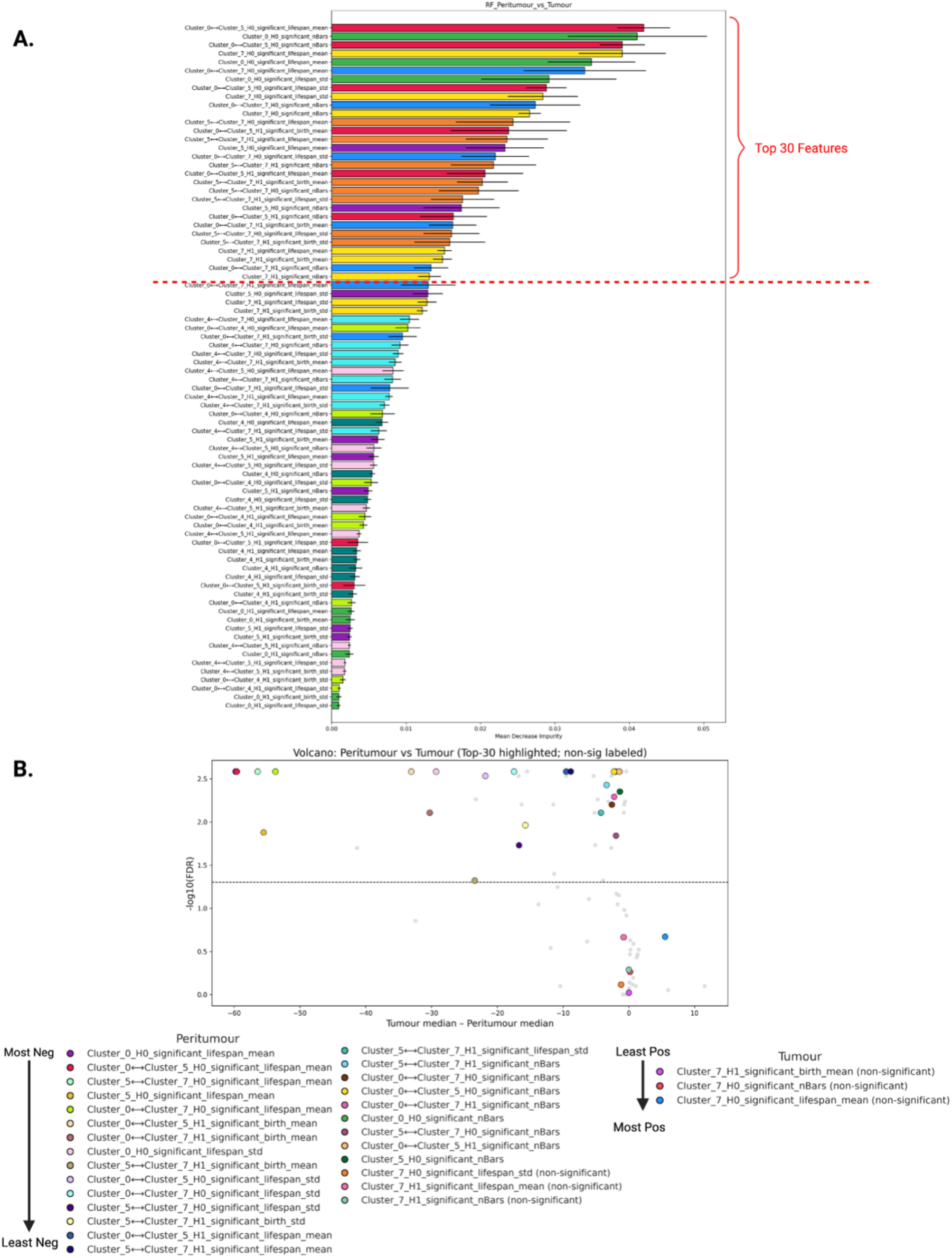
Feature selection for Cell DIVE TDA. **A)** Ranked bar plot of TDA features discriminating peritumour and tumour regions, as identified by random forest classifier feature importance (mean decrease in impurity, MDI). Each bar corresponds to a single TDA feature, derived from persistent homology calculations applied to single and pairwise neighbourhood clusters (e.g., NC0, NC4, NC5, NC7, and their pairings), and represents its mean importance across cross-validation folds. The top 30 features, as ranked by average MDI, are indicated above a dashed red line. **B)** Volcano plot comparing the distribution of the same TDA features between regions. For each feature, the x-axis gives the difference in slide-level medians (tumour minus peritumour). Y-axis is -log10 of the FDR-adjusted p-value from a paired Wilcoxon signed-rank test. Each point represents one TDA feature, with the top 30 features highlighted and labelled, and those not passing FDR significance are noted as non-significant in the legend.

**Supplementary Figure 10.**
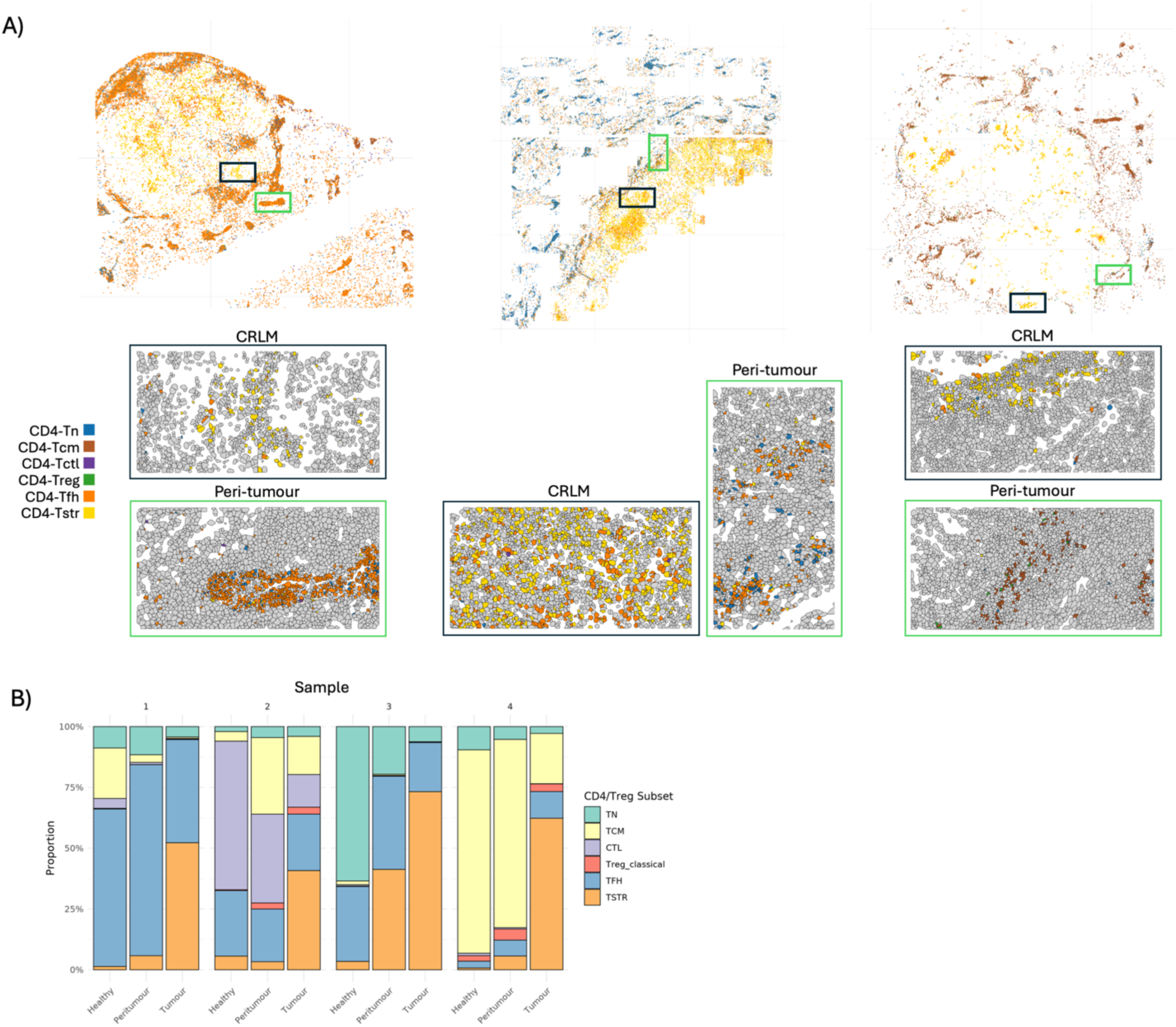
T-cell exhaustion and phenotypic states across tissues. **A)** Spatial maps of the 6 CD4 T-cell subsets. High-magnification regions within the CRLM and peri-tumoural liver demonstrate the differences in CD4 T-cell subsets in these locations, with the CD4 Tstr subset identified within the CRLM shown at higher magnification below through cell segmentation masks. **B)** Distribution of the indicated CD4 t-cell subtypes across the 4 specimens and within the 3 different tissue regions.

## References

1. Siegel, R. L., Wagle, N. S., Cercek, A., Smith, R. A. & Jemal, A. Colorectal cancer statistics, 2023. CA: A Cancer Journal for Clinicians 73, 233–254 (2023).

2. Brodt, P. Role of the Microenvironment in Liver Metastasis: From Pre- to Prometastatic Niches. Clin Cancer Res 22, 5971–5982 (2016).

3. Burkhardt, D. B., San Juan, B. P., Lock, J. G., Krishnaswamy, S. & Chaffer, C. L. Mapping Phenotypic Plasticity upon the Cancer Cell State Landscape Using Manifold Learning. Cancer Discov 12, 1847–1859 (2022).

4. Gui, P. & Bivona, T. G. Evolution of metastasis: new tools and insights. Trends in Cancer 8, 98–109 (2022).

5. Tufail, M., Jiang, C.-H. & Li, N. Immune evasion in cancer: mechanisms and cutting-edge therapeutic approaches. Sig Transduct Target Ther 10, 227 (2025).

6. Pancione, M., et al. Immune Escape Mechanisms in Colorectal Cancer Pathogenesis and Liver Metastasis. J Immunol Res 2014, 686879 (2014).

7. Che, L.-H., et al. A single-cell atlas of liver metastases of colorectal cancer reveals reprogramming of the tumor microenvironment in response to preoperative chemotherapy. Cell Discov 7, 80 (2021).

8. Wu, Y., et al. Spatiotemporal Immune Landscape of Colorectal Cancer Liver Metastasis at Single-Cell Level. Cancer Discov 12, 134–153 (2022).

9. Liu, Y., et al. Immune phenotypic linkage between colorectal cancer and liver metastasis. Cancer Cell 40, 424–437.e5 (2022).

10. Li, R., et al. Single-cell transcriptomic analysis deciphers heterogenous cancer stem-like cells in colorectal cancer and their organ-specific metastasis. Gut 73, 470–484 (2024).

11. Zhan, Y., et al. Single-cell transcriptomics reveals intratumor heterogeneity and the potential roles of cancer stem cells and myCAFs in colorectal cancer liver metastasis and recurrence. Cancer Letters 612, 217452 (2025).

12. Wang, F., et al. Single-cell and spatial transcriptome analysis reveals the cellular heterogeneity of liver metastatic colorectal cancer. Sci Adv 9, eadf5464 (2023).

13. Schürch, C. M., et al. Coordinated Cellular Neighborhoods Orchestrate Antitumoral Immunity at the Colorectal Cancer Invasive Front. Cell 182, 1341–1359.e19 (2020).

14. Vasquez, E. G., et al. Dynamic and adaptive cancer stem cell population admixture in colorectal neoplasia. Cell Stem Cell 29, 1213–1228.e8 (2022).

15. Gao, Y., et al. Spatial multi-omics reveals the potential involvement of SPP1+ fibroblasts in determining metabolic heterogeneity and promoting metastatic growth of colorectal cancer liver metastasis. Mol Ther 33, 3680–3700 (2025).

16. Zhang, Y., et al. CCL19-producing fibroblasts promote tertiary lymphoid structure formation enhancing anti-tumor IgG response in colorectal cancer liver metastasis. Cancer Cell 42, 1370–1385.e9 (2024).

17. Trebo, M., et al. Ex vivo modelling of human colorectal cancer liver metastasis by normothermic machine perfusion. Mol Cancer 24, 264 (2025).

18. Brown, L. V., Gaffney, E. A., Ager, A., Wagg, J. & Coles, M. C. Quantifying the limits of CAR T-cell delivery in mice and men. Journal of The Royal Society Interface 18, 20201013 (2021).

19. Xiao, Z., et al. Desmoplastic stroma restricts T cell extravasation and mediates immune exclusion and immunosuppression in solid tumors. Nat Commun 14, 5110 (2023).

20. CosMx® SMI Datasets. NanoString https://nanostring.com/products/cosmx-spatial-molecular-imager/Wpe-dataset/ (2025).

21. Meylan, M., et al. Tertiary lymphoid structures generate and propagate anti-tumor antibody-producing plasma cells in renal cell cancer. Immunity 55, 527–541.e5 (2022).

22. Wang, Q., et al. Tertiary lymphoid structures predict survival and response to neoadjuvant therapy in locally advanced rectal cancer. NPJ Precis Oncol 8, 61 (2024).

23. Le Rochais, M., et al. A Tertiary lymphoid structures-based pathological score predicts survival and recurrence in colorectal Cancer patients. Immunobiology 230, 152911 (2025).

24. Lazaris, A., et al. Vascularization of colorectal carcinoma liver metastasis: insight into stratification of patients for anti-angiogenic therapies. J Pathol Clin Res 4, 184–192 (2018).

25. Frentzas, S., et al. Vessel co-option mediates resistance to anti-angiogenic therapy in liver metastases. Nat Med 22, 1294–1302 (2016).

26. Immunological functions of liver sinusoidal endothelial cells - PubMed. https://pubmed.ncbi.nlm.nih.gov/27041636/.

27. Mukwaya, A., Jensen, L. & Lagali, N. Relapse of pathological angiogenesis: functional role of the basement membrane and potential treatment strategies. Exp Mol Med 53, 189–201 (2021).

28. Gajjala, P. R., et al. Dysregulated overexpression of Sox9 induces fibroblast activation in pulmonary fibrosis. JCI Insight 6, e152503 (2021).

29. Xie, Q., et al. High-Mobility Group A1 Promotes Cardiac Fibrosis by Upregulating FOXO1 in Fibroblasts. Front. Cell Dev. Biol. 9, (2021).

30. Li, W., et al. High-mobility group box 1 accelerates lipopolysaccharide-induced lung fibroblast proliferation in vitro: involvement of the NF-κB signaling pathway. Lab Invest 95, 635–647 (2015).

31. Cui, L., et al. Activation of JUN in fibroblasts promotes pro-fibrotic programme and modulates protective immunity. Nat Commun 11, 2795 (2020).

32. Zhao, L., et al. Tertiary lymphoid structures in diseases: immune mechanisms and therapeutic advances. Sig Transduct Target Ther 9, 225 (2024).

33. Maru, S. Y. et al. Antigen-presenting cancer-associated fibroblasts in murine pancreatic tumors differentially control regulatory T cell phenotype and function via CXCL9 and CCL22. bioRxiv 2025.03.27.645833 (2025) doi:10.1101/2025.03.27.645833.

34. Elyada, E., et al. Cross-Species Single-Cell Analysis of Pancreatic Ductal Adenocarcinoma Reveals Antigen-Presenting Cancer-Associated Fibroblasts. Cancer Discov 9, 1102–1123 (2019).

35. Zhang, Y., et al. Tertiary lymphoid structural heterogeneity determines tumour immunity and prospects for clinical application. Molecular Cancer 23, 75 (2024).

36. Muzio, L., Viotti, A. & Martino, G. Microglia in Neuroinflammation and Neurodegeneration: From Understanding to Therapy. Front. Neurosci. 15, (2021).

37. Wu, Z., et al. Peripheral nervous system microglia-like cells regulate neuronal soma size throughout evolution. Cell 188, 2159–2174.e15 (2025).

38. Wang, P. L., et al. Peripheral nerve resident macrophages share tissue-specific programming and features of activated microglia. Nat Commun 11, 2552 (2020).

39. Xing, L., et al. The role of neuropeptides in cutaneous wound healing: a focus on mechanisms and neuropeptide-derived treatments. Front. Bioeng. Biotechnol. 12, (2024).

40. Martier, A. T., et al. Estradiol and dihydrotestosterone exert sex-specific effects on human fibroblast and endothelial proliferation, bioenergetics, and vasculogenesis. Commun Biol 8, 1422 (2025).

41. Jin, S., Plikus, M. V. & Nie, Q. CellChat for systematic analysis of cell–cell communication from single-cell transcriptomics. Nat Protoc 20, 180–219 (2025).

42. Maud, M., et al. Memory CD4 T cells orchestrate neoadjuvant-responsive niches in colorectal cancer liver metastases. 2025.08.22.671704 Preprint at 10.1101/2025.08.22.671704 (2025).

43. Fernández Moro, C., et al. An idiosyncratic zonated stroma encapsulates desmoplastic liver metastases and originates from injured liver. Nat Commun 14, 5024 (2023).

44. Benjamini, Y. & Hochberg, Y. Controlling the False Discovery Rate: A Practical and Powerful Approach to Multiple Testing. Journal of the Royal Statistical Society: Series B (Methodological) 57, 289–300 (1995).

45. Zheng, L., et al. Pan-cancer single-cell landscape of tumor-infiltrating T cells. Science https://doi.org/10.1126/science.abe6474 (2021) doi:10.1126/science.abe6474.

46. Chu, Y., et al. Pan-cancer T cell atlas links a cellular stress response state to immunotherapy resistance. Nat Med 29, 1550–1562 (2023).

47. Asada, N., et al. The integrated stress response pathway controls cytokine production in tissue-resident memory CD4+ T cells. Nat Immunol 26, 557–566 (2025).

48. Lowery, F. J., et al. Neoantigen-specific tumor-infiltrating lymphocytes in gastrointestinal cancers: a phase 2 trial. Nat Med 31, 1994–2003 (2025).

49. Muscate, F., Woestemeier, A. & Gagliani, N. Functional heterogeneity of CD4+ T cells in liver inflammation. Semin Immunopathol 43, 549–561 (2021).

50. Lin, W.-P., Li, H. & Sun, Z.-J. T cell exhaustion initiates tertiary lymphoid structures and turbocharges cancer-immunity cycle. eBioMedicine 104, 105154 (2024).

51. Lowery, F. J., et al. Molecular signatures of antitumor neoantigen-reactive T cells from metastatic human cancers. Science 375, 877–884 (2022).

52. Janssen, E. M., et al. CD4+ T-cell help controls CD8+ T-cell memory via TRAIL-mediated activation-induced cell death. Nature 434, 88–93 (2005).

53. Lacher, S. B., et al. PGE2 limits effector expansion of tumour-infiltrating stem-like CD8+ T cells. Nature 629, 417–425 (2024).

54. Morotti, M., et al. PGE2 inhibits TIL expansion by disrupting IL-2 signalling and mitochondrial function. Nature 629, 426–434 (2024).

55. Polese, B., et al. Prostaglandin E2 amplifies IL-17 production by γδ T cells during barrier inflammation. Cell Reports 36, 109456 (2021).

56. Zheng, C., et al. Landscape of Infiltrating T Cells in Liver Cancer Revealed by Single-Cell Sequencing. Cell 169, 1342–1356.e16 (2017).

57. Guo, X., et al. Global characterization of T cells in non-small-cell lung cancer by single-cell sequencing. Nat Med 24, 978–985 (2018).

58. Zhang, L., et al. Lineage tracking reveals dynamic relationships of T cells in colorectal cancer. Nature 564, 268–272 (2018).

59. Di Giovannantonio, M., et al. Defining hypoxia in cancer: A landmark evaluation of hypoxia gene expression signatures. Cell Genomics 5, 100764 (2025).

60. Muhlich, J. L., et al. Stitching and registering highly multiplexed whole-slide images of tissues and tumors using ASHLAR. Bioinformatics 38, 4613–4621 (2022).

61. Schapiro, D., et al. MCMICRO: a scalable, modular image-processing pipeline for multiplexed tissue imaging. Nat Methods 19, 311–315 (2022).

62. Greenwald, N. F., et al. Whole-cell segmentation of tissue images with human-level performance using large-scale data annotation and deep learning. Nat Biotechnol 40, 555–565 (2022).

63. Liu, C. C., et al. Robust phenotyping of highly multiplexed tissue imaging data using pixel-level clustering. Nat Commun 14, 4618 (2023).

64. Nirmal, A. J. & Sorger, P. K. SCIMAP: A Python Toolkit for Integrated Spatial Analysis of Multiplexed Imaging Data. J Open Source Softw 9, 6604 (2024).

65. He, S., et al. High-plex imaging of RNA and proteins at subcellular resolution in fixed tissue by spatial molecular imaging. Nat Biotechnol 40, 1794–1806 (2022).

66. Stringer, C., Wang, T., Michaelos, M. & Pachitariu, M. Cellpose: a generalist algorithm for cellular segmentation. Nat Methods 18, 100–106 (2021).

67. Korsunsky, I., et al. Fast, sensitive and accurate integration of single-cell data with Harmony. Nat Methods 16, 1289–1296 (2019).

68. Danaher, P., et al. Insitutype: likelihood-based cell typing for single cell spatial transcriptomics. 2022.10.19.512902 Preprint at 10.1101/2022.10.19.512902 (2022).

69. Stuart, T., et al. Comprehensive Integration of Single-Cell Data. Cell 177, 1888–1902.e21 (2019).

70. Wood, S. N. Fast stable restricted maximum likelihood and marginal likelihood estimation of semiparametric generalized linear models. Journal of the Royal Statistical Society: Series B (Statistical Methodology) 73, 3–36 (2011).

71. Sathe, A., et al. Colorectal Cancer Metastases in the Liver Establish Immunosuppressive Spatial Networking between Tumor-Associated SPP1+ Macrophages and Fibroblasts. Clin Cancer Res 29, 244–260 (2023).

72. Fleischer, J. R., et al. Molecular differences of angiogenic versus vessel co-opting colorectal cancer liver metastases at single-cell resolution. Molecular Cancer 22, 17 (2023).

73. Pedregosa, F., et al. Scikit-learn: Machine Learning in Python. Preprint at 10.48550/arXiv.1201.0490 (2018).

74. Johnson, W. E., Li, C. & Rabinovic, A. Adjusting batch effects in microarray expression data using empirical Bayes methods. Biostatistics 8, 118–127 (2007).

75. Wolf, F. A., Angerer, P. & Theis, F. J. SCANPY: large-scale single-cell gene expression data analysis. Genome Biology 19, 15 (2018).

76. McInnes, L., Healy, J. & Melville, J. UMAP: Uniform Manifold Approximation and Projection for Dimension Reduction. Preprint at 10.48550/arXiv.1802.03426 (2020).

77. Becht, E., et al. Dimensionality reduction for visualizing single-cell data using UMAP. Nat Biotechnol https://doi.org/10.1038/nbt.4314 (2018) doi:10.1038/nbt.4314.

78. Bull, J. A., et al. MuSpAn: A Toolbox for Multiscale Spatial Analysis. 2024.12.06.627195 Preprint at 10.1101/2024.12.06.627195 (2025).

79. Otter, N., Porter, M. A., Tillmann, U., Grindrod, P. & Harrington, H. A. A roadmap for the computation of persistent homology. EPJ Data Sci. 6, 1–38 (2017).

80. Ali, D., et al. A Survey of Vectorization Methods in Topological Data Analysis. IEEE Transactions on Pattern Analysis and Machine Intelligence 45, 14069–14080 (2023).

81. Gregorutti, B., Michel, B. & Saint-Pierre, P. Correlation and variable importance in random forests. Stat Comput 27, 659–678 (2017).

82. Breiman, L. Random Forests. Machine Learning 45, 5–32 (2001).

83. Virtanen, P., et al. SciPy 1.0: fundamental algorithms for scientific computing in Python. Nat Methods 17, 261–272 (2020).

